# Identification of PUFA interaction site on ASIC3 suggests a complex regulatory mechanism

**DOI:** 10.64898/2026.01.02.697424

**Authors:** Rebecca Roth, Ramya Bandarupalli, Robert C. Klipp, Jing Li, John R. Bankston

**Author notes:** These authors contributed equally to this work (Rebecca Roth, Ramya Bandarupalli). Corresponding Author: John Bankston, Department of Physiology and Biophysics, Research Complex 1 North, University of Colorado Anschutz Medical Campus, Aurora, CO, 80045.; Jing Li.

## Abstract

Inflammatory mediators including polyunsaturated fatty acids (PUFAs) potentiate ASIC3 by altering channel gating, yet the molecular basis for their interactions remains poorly defined. Using all-atom MD simulations and electrophysiology, we show that DHA accumulates around ASIC3 through interactions with a membrane-facing electropositive region along the outer leaflet of TM1. In the open state, a more buried binding site for the lipid is exposed that allows the carboxylate head group to bind a critical arginine along with nearby polar residues. Mutation of this site eliminates the DHA-mediated slowing of channel desensitization and also modestly reduces the effect PUFAs have on the proton sensitivity of channel activation. The same mutations do not prevent regulation by other lipid classes like N-acyl amino acids or lysophosphatidyl choline. Our results provide the first detailed description of a PUFA binding site on ASICs, offering new insights into lipid modulation and potential strategies for developing novel treatments for inflammatory pain.

## INTRODUCTION

Cell membranes are composed of a wide diversity of lipids that can have profound impacts on the activity of membrane proteins. There is a long history describing the regulation of ion channel function by the lipid environment but mechanisms of modulation are complex and vary between different ion channels^1–6^. In general, channel regulation is thought to occur through two major mechanisms: indirect effects on conformational rearrangements through changes in the lipid bilayer (i.e. thickness, fluidity, curvature), or alterations to channel gating through stable interactions. Characterizing the interactions between lipids and ion channels has become even more complex^7–9^. Several hypotheses have been presented including stable binding of lipids to a binding site on the channel, non-selective transmembrane domain solvation, and/or the formation of local lipid ring (annulus) around the channel^10–13^. Understanding these dynamic interactions remains a critical challenge in the field, highlighting the need for both experimentation and computational approaches to disentangle their relative contributions influencing channel gating and modulation.

Over the last 20 years, there has been increasing evidence that Acid-Sensing Ion Channels (ASICs) are critically modulated by several lipid classes^3,14–16^. ASICs are pH-activated, sodium-permeable ion channels that belong to the Deg/ENaC family of ion channels and are highly expressed in the central (CNS) and peripheral (PNS) nervous systems where they canonically serve as pH sensors^17,18^. The ASIC3 isoform is the most sensitive to proton-mediated activation and is predominantly expressed in nociceptive sensory neurons^19,20^. Inhibiting ASIC3 activity through pharmacological blockade or genetic deletion has been shown to prevent the development of chronic muscle and tactile hyperalgesia in response to repeated injections of acid, formalin, reserpine, and carrageenan^21–24^. Proinflammatory mediators released during inflammation including neurotrophins (i.e. NGF), cytokines, serotonin, and lipids, have been shown to increase sensory neuronal excitability through modulation of ASICs^14,22,24^. Work from our lab and others have identified at least three classes of single acyl chain lipids that can potentiate ASIC3 currents: polyunsaturated fatty acids (PUFAs), lysophosphatidylcholines (LPCs), and N-acyl amino acids (NAAAs)^3,14,15^. While structurally distinct, all three lipid classes exert qualitatively similar effects on the channel although NAAAs and LPCs seem to cause larger changes in channel function. Each lipid slows the rate of channel desensitization, shifts the pH dependence of activation in the basic direction, and increases the current even at saturating proton concentrations. A careful examination of the structural requirements for lipid regulation of ASIC3 suggested roles for both the head and tail group of the lipid^3^. However, the electronegativity of the head group demonstrated to be the most critical element in determining the strength of potentiation.

PUFA concentrations increase during many pathological conditions such as ischemia, epilepsy, tissue injury, and inflammation^25–27^. Their release from phospholipids via activation of phospholipases during these events can allow PUFAs to move freely through cells to influence signaling pathways that regulate gene expression, receptor signaling, and membrane dynamics^28^. Additionally, PUFAs can be metabolized into analogs which can either have similar or opposing effects on ion channel regulation in comparison to their precursors, depending on their new structural properties.

Marra and colleagues demonstrated that inflammatory exudates with neutral pHs from human patients were also sufficient to activate ASICs including ASIC3^14^. The active components of the exudates were found to be two different classes of single acyl chain lipids that were present at high concentrations: an ω-6 PUFA known as arachidonic acid (AA) and LPC. The combination of these lipids was sufficient to activate ASIC3 in the absence of acidification, and injections of LPC and AA into mouse hind paws produced significant pain behaviors that were reduced in ASIC3 KO mice or mice co-injected with an ASIC3-specific inhibitor APETx2. This provides evidence that ASIC3 may be a viable target for pharmacological inhibition in inflammatory pain, which requires a better understanding of the interactions between these lipids and the channel.

Work from our lab and others has pointed to a region on TM1 near the outer leaflet as potentially being critical for lipid regulation of ASIC^3^. However, these studies provide an incomplete picture. We previously mutated several residues in this region and showed a reduced effect of the PUFA docosahexaenoic acid (DHA) but did not have evidence of other potential interactions. Molecular dynamics (MD) simulations on human ASIC3 suggested a pair of arginine residues near the outer leaflet may be critical, but one of those arginine residues is not present in other isoforms and orthologues that are still sensitive to lipid modulation^29^. To overcome these limitations, we combined Molecular Dynamics and patch clamp electrophysiology to determine the residues on ASIC3 that may be critical for lipid interaction and functional regulation of the channel. We discovered several residues that are required for both interaction and the change in channel function. Overall, we hypothesize that the data we present here points to complex regulation of the channel by lipids and that a model where the lipid stably binds to the channel is likely insufficient to describe modulation of ASIC3 by single acyl chain lipids.

## Results

### DHA potentiates ASIC3 activation through preferential binding to the extracellular leaflet

DHA, the most abundant PUFA in the brain, acts as a strong potentiator of both ASIC1 and ASIC3 channels^3,15^. Structurally, DHA contains a carboxyl head group and a 22-carbon chain tail group which contains 6 cis double bonds throughout the acyl chain (Fig. 1A). Previous work on PUFA modulation of ASICs from our group and others has largely focused on the other abundant PUFA in the brain: arachidonic acid (AA). However, DHA exhibits much stronger potentiating effects on ASIC3 gating including a larger shift in the pH-dependence of activation and a larger slowing of channel desensitization. All PUFAs have the same carboxyl head group which we have shown is the primary driver of the potentiation, so we elected to focus this study on DHA as a model lipid for understanding, generally, how PUFAs bind to and modulate ASIC3 channels.

**Figure 1.**
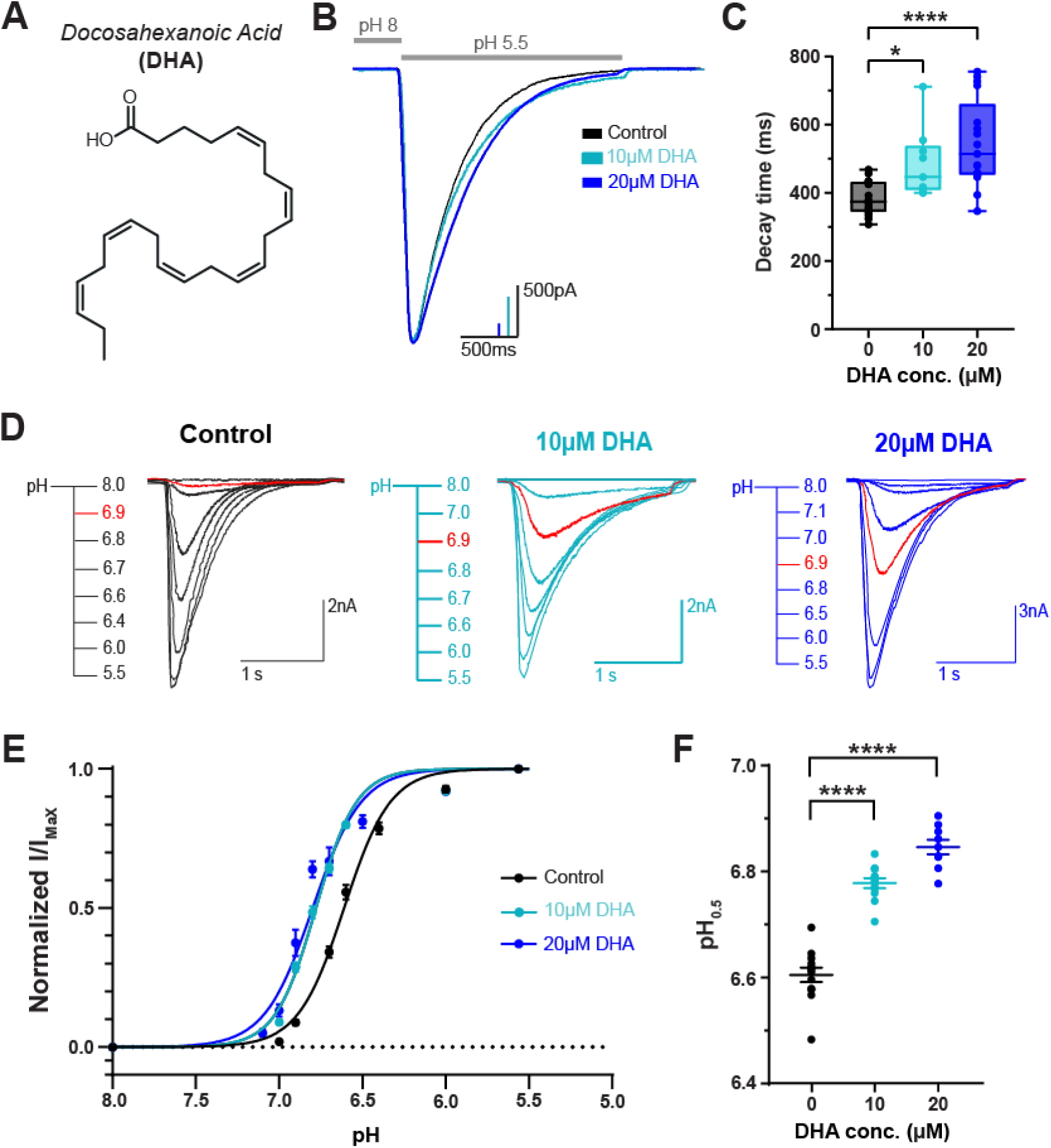
DHA potentiates ASIC3 gating effects in a dose-dependent manner. **(A)** Chemical structure of DHA, a carboxylic acid with a 22-carbon chain and 6 cis double bonds starting from the 3^rd^ carbon. **(B)** Representative whole-cell recordings showing the slowing of the desensitization rate of ASIC3 ± 10 and 20 µM DHA. Currents were elicited by performing fast-switch perfusion changes between pH 8 and pH 5.5 solutions. **(C)** Box plot showing the time point at which ASIC3 currents decayed from their peak by 63% (1/e) at 0, 10 and 20 µM DHA (0μM: 385.5 ± 10.9 ms, *n = 19*; 10 μM: 485.7 ± 33.9 ms, *n = 9*; 20 μM: 545.8 ± 30.7 ms, *n = 17*). **(D)** Representative whole-cell recordings showing pH-dependent activation of ASIC3 ± 10 and 20 µM DHA. Traces elicited at pH 6.9 are highlighted in red to visualize the increase in pH sensitivity in the presence of DHA concentrations. **(E)** Fitted curves showing the pH dependence of activation of ASIC3 WT at different concentrations of DHA, fitted using a Hill-type equation (see Equation 1 in Methods). **(F)** Average activation pH_0.5_ values for ASIC3 channels in response to 0, 10, and 20 µM DHA (0μM: pH_0.5_ = 6.60 ± 0.014, *n = 13*; 10 μM: pH_0.5_ = 6.78 ± 0.009, *n = 12*; 20 μM: pH_0.5_ = 6.85 ± 0.014, *n = 9*). All data given as mean ± SEM. C-D, One-way ANOVA post hoc Dunnett’s test. ***, P < 0.001; ****, P < 0.0001 (see Methods for details).

To confirm and extend our previous measurements of the modulatory effects of DHA on ASIC3 channels, we performed whole-cell patch-clamp recordings in Chinese hamster ovary (CHO) cells transfected with rat ASIC3 (rASIC3) and measured currents in the absence and presence of 10 or 20 µM DHA. To ensure the effect was saturated, cells were preincubated for at least 5 minutes in DHA-containing pH 8 solution prior to each recording. We previously showed that the effects of DHA were largely saturated in 1-2 minutes^3^. Consistent with our previous work, the application of DHA leads to multiple changes in channel function. First, addition of both 10 and 20 µM DHA resulted in a consistent and reproducible slowing of channel desensitization rate which can be seen when we overlay currents elicited by a fast solution switch from pH 8 to pH 5.5 (Fig. 1B). To quantify the slowing of channel desensitization, we measured the time it takes for the current to decay to 1/e (63%) of the peak value. Application of 10 and 20 µM slowed the rate of channel desensitization in a concentration-dependent manner, slowing the rate from 385 ms to 486 and 546 ms respectively (Fig. 1C, Supplementary Table 1). In addition to a change in desensitization rate, 10 and 20 µM DHA also increased the proton sensitivity of channel activation. To demonstrate this, we made solution switches from pH 8 to varying pH levels between 7.1 and 5.5 (Fig. 1D) and then plotted the peak current as a function of the activating pH (Fig. 1E). Consistent with our previous findings, ASIC3 currents show a half-maximally activating pH value (pH_0.5_) of 6.6 in the absence of lipid. In the presence of 10 µM and 20 µM DHA, the pH_0.5_ exhibited an alkaline shift by about 0.18 and 0.25 pH units respectively (Fig. 1F, Supplementary Table 1).

It has been hypothesized for many ion channels, including ASICs, that the negatively charged head group of PUFAs likely makes stable electrostatic interactions or hydrogen bonds with side chain atoms of arginine (R), lysine (K), and tyrosine (Y) residues located along their TM segments to alter channel gating^3,6,30,31^. Both TM1 and TM2 of ASIC3 have positively charged residues positioned near the inner and outer leaflet of the channel, indicating the possibility that there are interaction sites for lipids with negatively charged head groups on either or both sides of the channel. It has been shown that AA shifts the pH-dependence of ASIC2a channel activation upon application to either the intracellular or extracellular side of the channel, while we previously showed that a PUFA derivative of AA known as N-arachidonyl glycine (AG) only produces potentiating effects on ASIC3 when applied to the outer leaflet of the membrane^3,15^. Therefore, we asked whether PUFAs exhibited a sidedness for potentiating ASIC3 channels.

Here, we tested the hypothesis that DHA potentiates ASIC3 currents only when it is applied to the extracellular side of the channel. To do this, we applied 20 µM DHA to the patch pipette and elicited channel activation by applying brief external applications of a mildly activating pH 6.8 solution (Fig. 2A). We saw no changes in the peak current across several sweeps when compared to control conditions, whereas acute external application of 20µM DHA produced a 2.3-fold increase in current amplitude size within 45 seconds of exposure (Fig. 2B). It is possible that the time it takes to start the recording resulted in saturation of the effect prior to our measurement, thus, we also looked at changes in the steady state parameters. There were no significant changes to the pH sensitivity for channel activation or the rate in channel desensitization when DHA was internally applied for 5 or 10 minutes (Fig 2C-E). Altogether, this strongly suggests that DHA, like AG, alters ASIC3 channel desensitization and pH sensitivity through action on the extracellular side of the channel.

**Figure 2:**
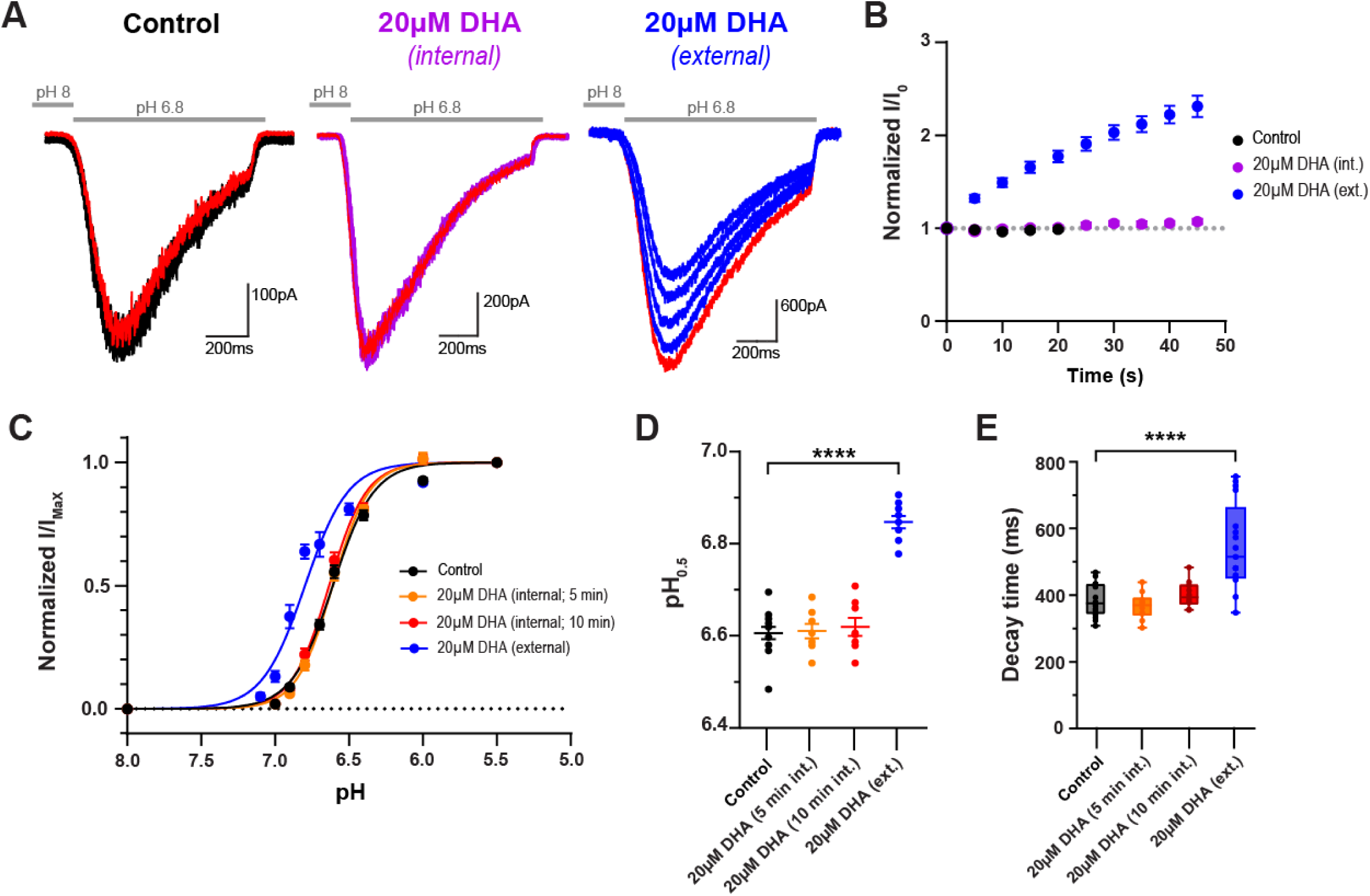
Exposure to the extracellular leaflet is necessary to induce DHA-mediated potentiation of ASIC3 currents. **(A)** Representative traces of the first 5 sweeps within a single cell showing repeated pH 6.8-evoked currents in the absence (right) or presence of either internal (middle) or external (left) application of 20 µM DHA. The red trace represents the current elicited during the 5^th^ sweep of the time course in B. **(B)** Time course of pH 6.8-evoked ASIC3 currents. Internal application represents 20 µM DHA within the patch pipette; external applications represent 20 µM DHA within the pH 8 bath solution (*n* = 7-10 cells/ condition). **(C)** Fitted curves showing the pH dependence of activation of ASIC3 WT following different applications of 20 µM DHA, fitted using a Hill-type equation (see Equation 1 in Methods). **(D)** Average activation pH_0.5_ values for ASIC3 channels in response to internal or external application of 20 µM DHA (<u>0μM:</u> pH_0.5_ = 6.60 ± 0.014, *n = 13;* <u>20 μM int. 5 min:</u> pH_0.5_ = 6.61 ± 0.016, *n = 8;* <u>20 μM int. 10 min:</u> pH_0.5_ = 6.62 ± 0.02, *n = 8;* <u>20 μM ext.:</u> pH_0.5_ = 6.85 ± 0.014, *n = 9*). **(E)** Box plot showing the time point at which ASIC3 currents decayed from their peak by 63% (1/e) in response to internal or external application of 20 µM DHA (<u>0</u> <u>μM:</u> 385.5 ± 10.9 ms, *n = 19*; <u>20 μM 5 min int.:</u> 366.8 ± 12.8 ms, *n = 10*; <u>20 μM 10 min int.:</u> 402.1 ± 12.6 ms, *n = 10*; <u>20 μM ext.:</u> 545.8 ± 30.7 ms, *n = 17*). C-E, ASIC3 WT and external 20 µM DHA are replotted from Fig. 1 for comparison. All data given as mean ± SEM. D-E, One-way ANOVA post hoc Dunnett’s test. ****, P < 0.0001 (see Methods for details).

### DHA accumulates near ASIC3, but channel opening exposes an arginine for stable binding

Work from our lab and others have suggested that a cluster of arginine residues located in the outer leaflet of TM1 are potentially critical for mediating lipid binding to ASIC1 and ASIC3^3,29^. However, previous MD simulations and electrophysiology data support differing conclusions about which outer leaflet residues are critical for functional regulation of the channel. Therefore, to better understand how lipid interactions with ASICs lead to changes in function, we employed all-atom MD simulations to determine, at atomic resolution, how DHA interacts with the TMs of ASIC3 in different conformational states. With the predictions from these models, we then performed site-directed mutagenesis and patch-clamp electrophysiology to understand the impact these residues have on lipid regulation of the channel.

To perform MD simulations, homology models of human ASIC3 (hASIC3) were generated based on the chicken ASIC1 structure in the open state (PDB ID 4NTY) and the human ASIC1 structure in the resting state (PDB ID 7CFS) as no structures of ASIC3 have been solved to date^32,33^. To assess the reliability of our homology model we compared it with both the open-state chicken ASIC1 structure and an AlphaFold prediction of human ASIC3 (AF-Q9UHC3-4). All three structures show strong overall agreeement in the backbone architecture and in the transmembrane regions. We demonstrated this by showing the positioning and stability of key sidechains were consistent across models as evidenced by stability of the χ1 dihedral angle of a key sidechain in TM1 as well as the stability of the secondary structure of TM1 throughout the simulation (Supplementary Figure 1).

To probe spontaneous lipid-protein interactions, DHA molecules were randomly distributed within the lipid bilayer at a 10:1 POPC: DHA molar ratio (Fig. 3, Table 1; see Methods). The systems were equilibrated and subsequently simulated for 5 μs using a single production trajectory to identify DHA-interaction hotspots on hASIC3. During these simulations, DHA molecules diffused within the lipid bilayer and began to accumulate at sites on the channel near both the inner and outer leaflet of the membrane. In the resting state, densities representing the head group of DHA can been seen accumulating at sites both on the intracellular and extracellular side of the channel (Fig. 3A-D). DHA clustering was most pronounced around a pair of adjacent positively charged residues arginine 65 and 68, termed hR65 and hR68 here to indicate they are residues from the human isoform, located in the outer leaflet of TM1, which appear to form a distinct electropositive region that attracted the negatively charged DHA headgroups (Fig. 3B,C). Additionally, the head group of DHA molecules also clustered near positively charged residues in the intracellular leaflet including hR41, hR42, hR465, and hK467 (Fig. 3B). Notably, DHA interactions with inner-leaflet residues were observed only in the resting-state trajectory, as the intracellular arginine residues were unresolved in the 4NTY structure used for homology modeling of the open state.

**Figure 3.**
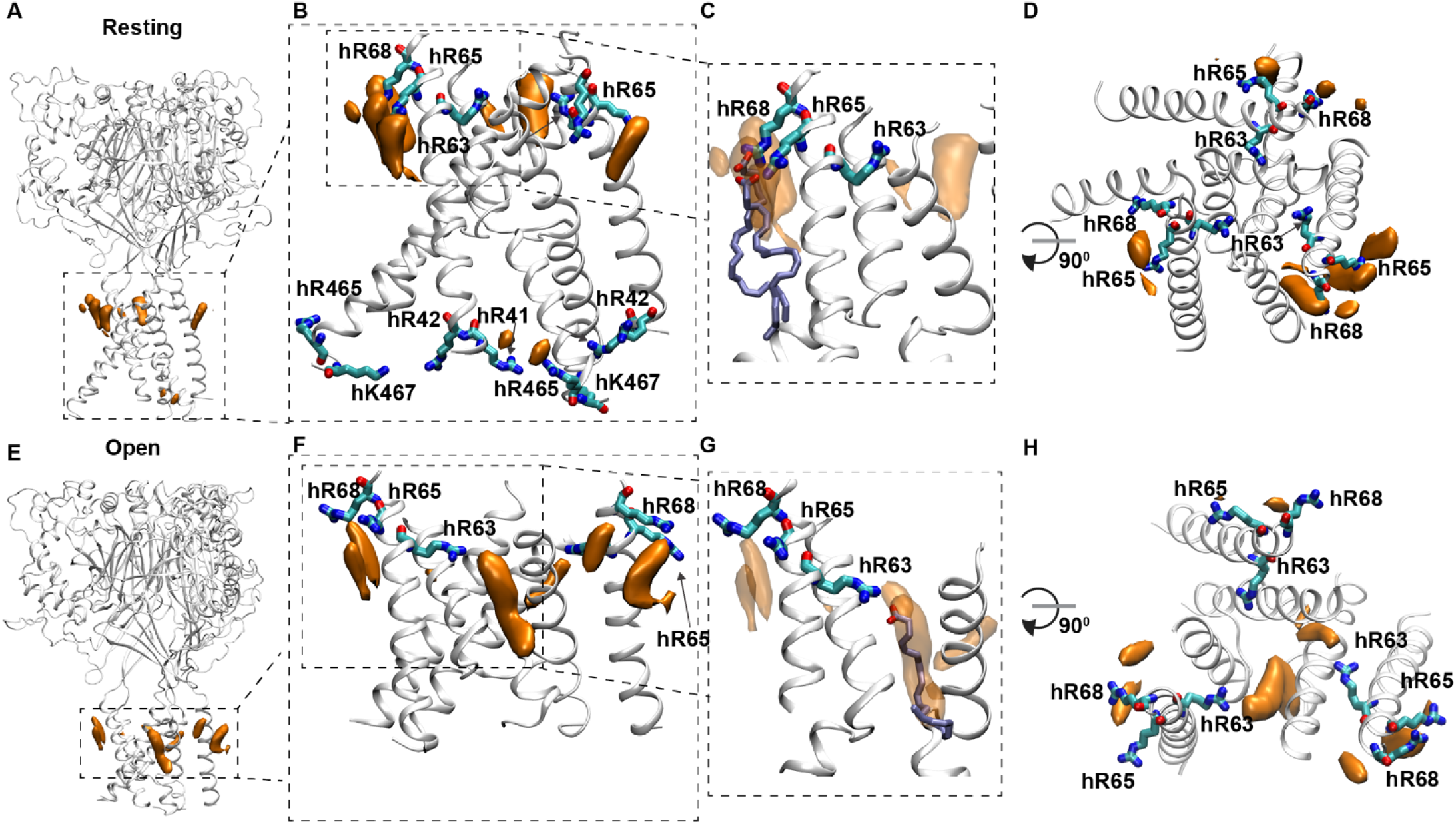
Spontaneous binding of DHA is more stable when ASIC3 is in the open state. **(A)** Side view of the full ASIC3 resting-state homology model. **(B)** Zoomed-in view of the TM region, **(C)** zoomed-in view of the arginine-rich cluster in the upper leaflet, and **(D)** top view of the resting-state hASIC3 model showing spontaneous accumulation of DHA around both the intracellular and extracellular ends of the TM helices. DHA is shown in an ice blue licorice representation (panel C), while orange isosurfaces represent DHA density (panel A-D). DHA preferentially localizes near the arginine-rich region formed by residues hR65 and hR68 (cyan), creating a lipid density in the outer leaflet of the membrane. Additional DHA densities are also observed towards the intracellular leaflet. **(E)** Side view of the full ASIC3 open-state homology model. **(F)** Zoomed-in view of the TM region, **(G)** zoomed-in view of the arginine-rich cluster, and **(H)** top view of the open-state hASIC3 model showing spontaneous accumulation of DHA primarily at the extracellular ends of the TM helices. Densities were calculated using VMD VolMap and represent the average distribution of DHA across the trajectory, (panels A–D, Trajectory 1, the spontaneous DHA-partitioning simulation of the resting-state hASIC3 model; panels E–H, Trajectory 2, the corresponding simulation of the open-state model; see Table 1 for a full description of all trajectories), visualized as isosurfaces at an isovalue of 0.4.

**Table 1:**
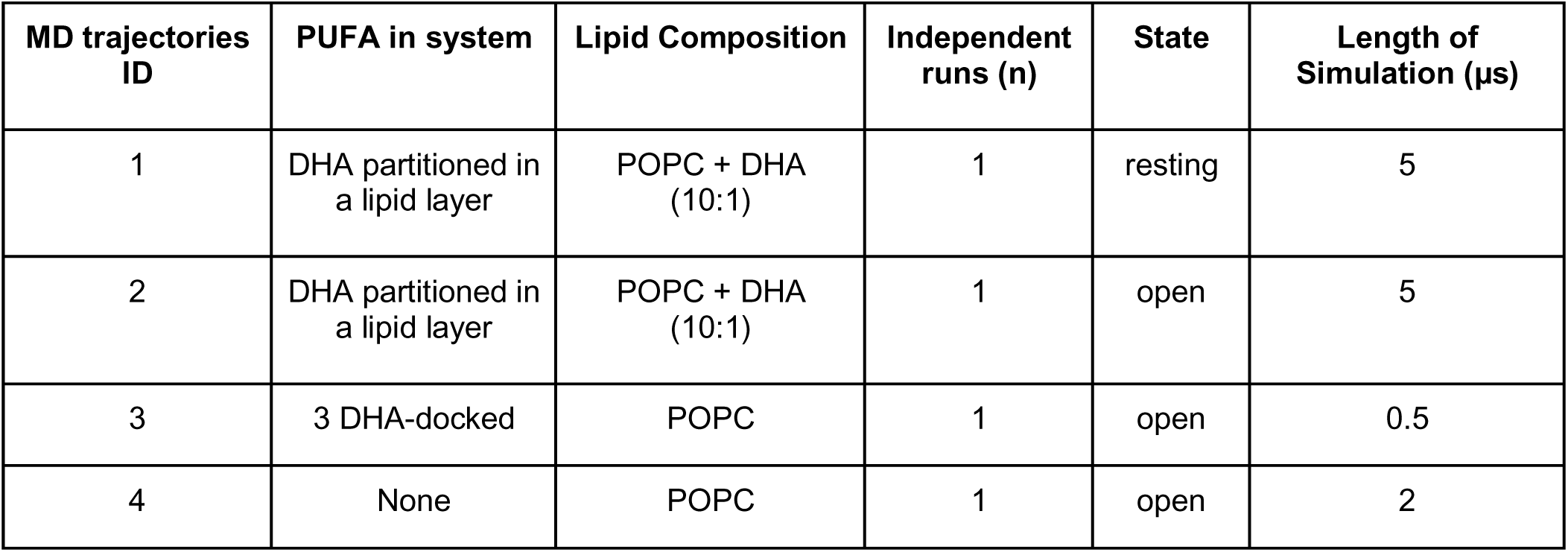
Summary of MD simulations performed throughout study.

| MD trajectories ID | PUFA in system | Lipid Composition | Independent runs (n) | State | Length of Simulation (μs) |
| --- | --- | --- | --- | --- | --- |
| 1 | DHA partitioned in a lipid layer | POPC + DHA (10:1) | 1 | resting | 5 |
| 2 | DHA partitioned in a lipid layer | POPC + DHA (10:1) | 1 | open | 5 |
| 3 | 3 DHA-docked | POPC | 1 | open | 0.5 |
| 4 | None | POPC | 1 | open | 2 |

In the open state, a large density absent in the resting state appeared near residue hR63 (Fig. 3E-H). DHA molecules continued to interact with hR65 and hR68, but accumulated much more densely around hR63, forming a distinct higher-density lipid cluster at the outer membrane interface (Fig. 3F,G). This difference is likely due to its accessibility; in the resting state, hR63 is positioned deep within the protein interior, shielded from the surrounding lipid bilayer. When the channel opens, TM1 moves outward, repositioning hR63 toward the membrane interface and exposing its positively charged side chain to the lipid environment. This structural rearrangement creates a new electropositive binding surface that is more buried in the channel interior that allows the DHA headgroup to engage with hR63 directly through stable electrostatic interactions.

To look more quantitatively at the interaction interface for DHA on the extracellular side of the channel, we calculated time-averaged occupancy maps which reflect the fraction of time that the head group of DHA molecules occupied a particular position in space relative to each individual residue (Fig. 4A-C). These heat maps show an increased concentration of DHA molecules surrounding the channel centered at these arginine residues. In addition, using a distance vs. time plot showing the proximity of the carboxylate carbon of DHA and the terminal carbon to the side chain (Supplementary Figures 2-4), we calculated the DHA contact duration at residues hR63, hR65, and hR68 in both states. In both the open and resting state simulations, DHA molecules were recruited to this region throughout the 5µs simulations, yet their distributions differed markedly between two states. In the resting state, DHA primarily interacted with hR65 and hR68, exhibiting mean lifetimes of 162 and 124 ns, respectively, and the longest individual binding events lasting up to 1.2 and 2.9 µs (Fig. 4D). DHA density near hR63 was minimal in the resting conformation and no detectable binding events were observed during the 5 µs trajectory, reflecting its buried position (Fig. 4B, D, Supplementary Figure 5).

**Figure 4.**
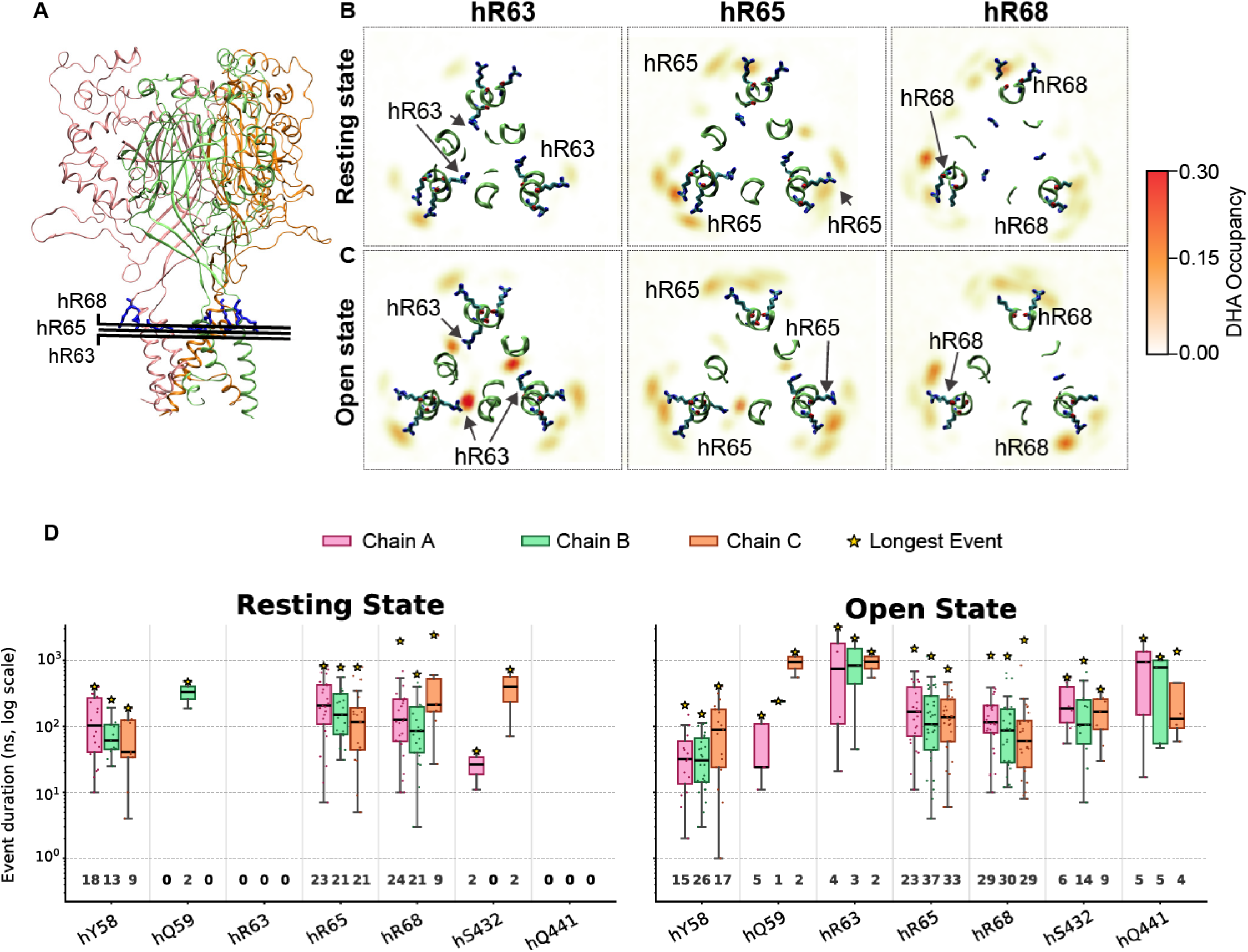
Mutation of R63 eliminates the effect of DHA on channel desensitization. **(A)** Cross-sectional view of the ASIC3 trimer highlighting the levels used to generate the occupancy maps in panels B and C. The protein is shown in ribbons with three subunits colored pink, orange, and green. 2D volume maps illustrate the DHA occupancy near residues R63, R65, and R68 in the resting **(B)** and open **(C)** state simulations (Trajectory 1, resting state; Trajectory 2, open state; see Table 1). The color scale represents the fractional occupancy of DHA, where a value of 0.3 corresponds to 30% occupancy during the simulation. Yellow to red regions indicate higher DHA density*. (***D**) Box-and-whisker plots showing the distribution of individual DHA binding event durations (ns, log scale) at seven residues across three subunits in the open state (left) and resting state (right). Residues are labeled by chain identity (Chain A, pink; Chain B, green; Chain C, orange) for reference; however, DHA binding occurs at inter-subunit interfaces (A–B, B–C, and C–A) rather than within a single subunit. A binding event was defined as a continuous period during which the smoothed minimum distance between the DHA carboxylate carbon and the target residue fell below 4.3 Å. Each box shows the interquartile range with the median indicated by the central line; whiskers extend to 1.5× IQR; individual events are overlaid as jittered dots. The gold star (★) marks the longest binding event within each group. The number of events (*n*) is shown below each box.

In contrast, in the open state of the channel a binding surface deeper within the channel interior is exposed. This is highlighted by the formation of a stable interaction between hR63 and DHA, characterized by fewer total events but much longer durations. The mean duration of these binding events were about 495 ns, with the longest individual binding event lasting over 3 µs (Fig. 4D). Additionally, multiple detectable binding events were made at hQ59 and hS432 with randomly partitioned DHA molecules throughout both open and resting state simulations however the frequency of those interactions increased in the open state. Finally, interactions were only detectable at hQ441 in the open state. Meanwhile, hR65 and hR68 continued to form dynamic, recurrent interactions, with mean lifetimes of 142 and 113 ns (Fig. 4D). Interestingly, the occupancy at hR63 in the open state exceeded that observed at hR65 and hR68 in either conformational state (Fig. 4D). This suggests that a more surface interaction site for the lipid exists regardless of channel state, but a longer-lived and more stable binding site, comprised partly of hR63, hQ59, hS432, and hQ441 is exposed upon channel opening.

### Molecular docking reveals other key residues which mediate effects on desensitization

The spontaneous binding simulations showed that randomly inserted DHA molecules diffuse through the bilayer and accumulate near the extracellular end of TM1, forming long-lived interactions with a cluster of arginine residues. Upon channel opening, a more buried binding cite is exposed leading to very stable interaction with hR63 in the open state. To further assess whether nearby residues contribute to DHA stabilization and to characterize the persistence of headgroup interactions, we performed molecular docking followed by all-atom MD simulations for 500 ns. In contrast to the random DHA partitioning approach, which allowed DHA to freely explore the membrane environment, molecular docking enables targeted placement of DHA at the three equivalent binding sites, on each of the ASIC3 subunits that comprise the functional channel, centered on hR63 and thereby allowing a systematic evaluation of DHA interactions with surrounding residues during extended MD simulations.

Contact analysis of the docked MD trajectories revealed that, in addition to the persistent interaction with hR63, the DHA headgroup frequently engaged in contacts with several nearby polar residues (Fig. 5A, Supplementary Figure 6). The contact number, defined as the number of atomic pairs between DHA and a residue that are within 4 Å in each simulation frame, was used to quantify the strength and persistence of these interactions over time (Fig. 5B). We examined the interactions between each of the three DHA molecules. For each subunit, hQ59 and hQ441 displayed average contact numbers greater than 1, indicating that multiple atomic pairs between these residues and the DHA carboxylate group remain within proximity. hS432 also interacted frequently with DHA, showing the average contact number as 0.73, which is significantly higher than other nearby residues such as hE433 and hD437. These observations are confirmed by looking at both the mean contact pairs and contact frequency and of the residues near this buried binding site (Fig. 5C, D). Taken together these data are consistent with a stable interaction in the open state with the more interior binding site.

**Figure 5.**
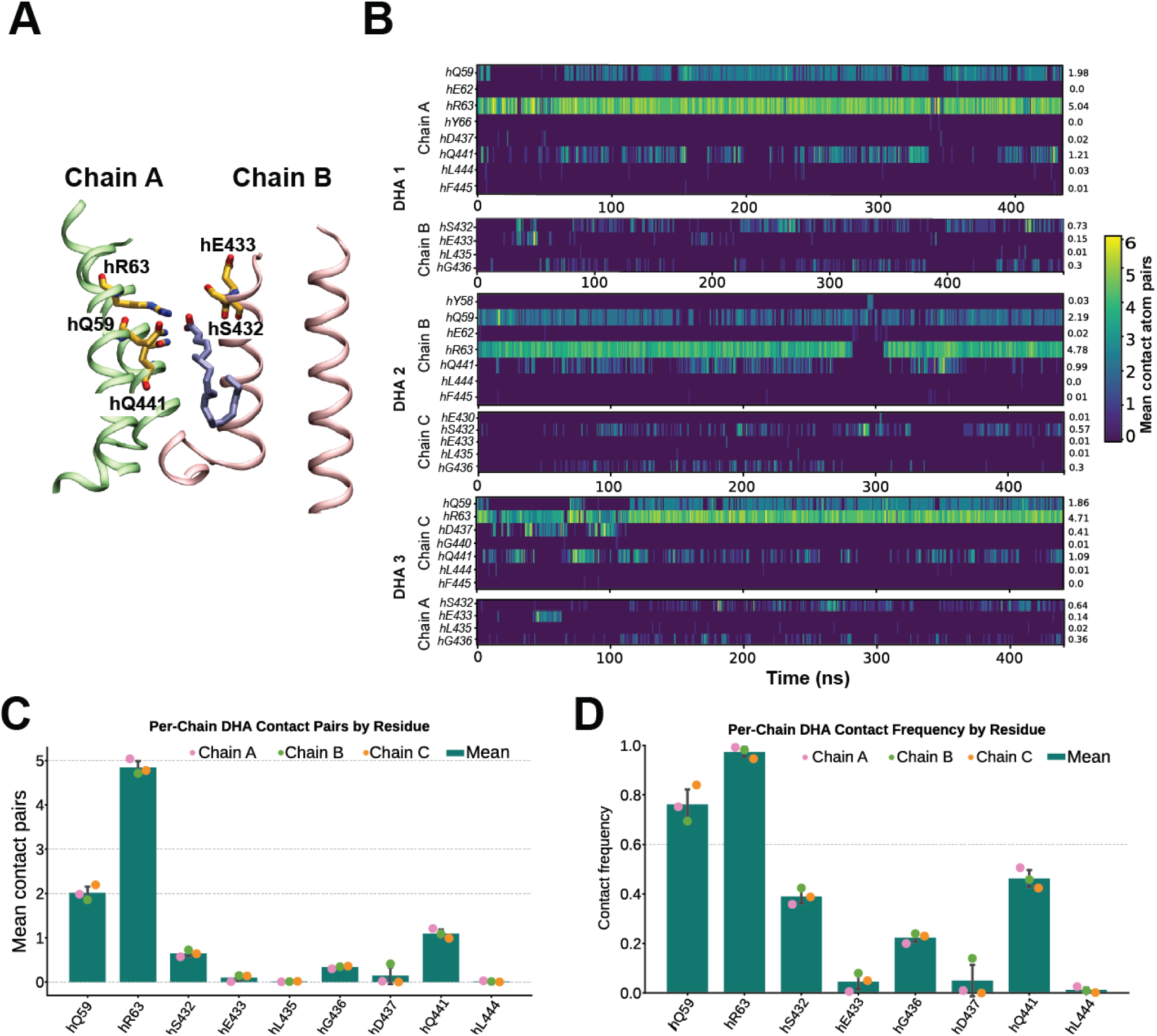
Other key residues nearby R63 contribute to PUFA interaction with ASIC3. **(A)** Molecular visualization from trajectory 3 (the 500 ns simulation of the open-state hASIC3 model with three DHA molecules pre-docked near the R63 binding site; see Table 1) showing DHA bound near the TM region. The protein is shown in ribbons with two chains colored lime and pink. DHA is displayed in ice-blue licorice, and the residues from both chains that interact with DHA are shown in orange licorice representation. **(B)** Time-resolved contact maps showing DHA–residue interactions for each DHA molecule (DHA 1–3) across the three ASIC3 subunits (Chains A–C). The x-axis represents simulation time (ns), and the y-axis lists residues. Color intensity reflects the number of heavy-atom pairs within a 4 Å cutoff between DHA and each residue at each frame. Values shown on the right indicate the mean contact number for each residue over the trajectory. (**C**) Bar plot of mean DHA– residue contact pairs averaged over time for each residue. Bars represent the mean across the three subunits, while individual points indicate values for each chain (Chain A, pink; Chain B, green; Chain C, orange). (**D**) Contact frequency (fraction of simulation frames with at least one DHA–residue contact within 4 Å) for each residue. Bars represent the mean across subunits, with individual points showing per-chain values.

### Mutation of a single arginine prevents DHA-mediated slowing in desensitization rate

To validate the predictions of our simulated models, we created mutations to residues identified as hotspots and measured their impacts on DHA-mediated slowing of channel desensitization. Given that DHA acts on ASIC3 currents only when applied to the extracellular side of the channel, we focused our mutagenesis only on the residues located near the outer leaflet of the membrane. Mutations were made using the rat ASIC3 gene to stay consistent with our previous work and to ensure optimal current magnitudes within our cell system because, in our hands, rat ASIC3 generates larger currents when compared to human ASIC3 and many of the mutations made along the TMDs tend to greatly reduce expression (Table 2). The numbering of rat and human ASIC3 does not align perfectly (Supplementary Figure 7). To ensure clarity for which position we are discussing we will continue to use the first letter of the orthologue (h for human and r for rat) along with the position of the residue in the relevant orthologue. For instance, when we discuss arginine 63, which is the equivalent of R64 in rat we will call them hR63 and rR64, respectively, or when relevant show the residue for both orthologues.

**Table 2:**
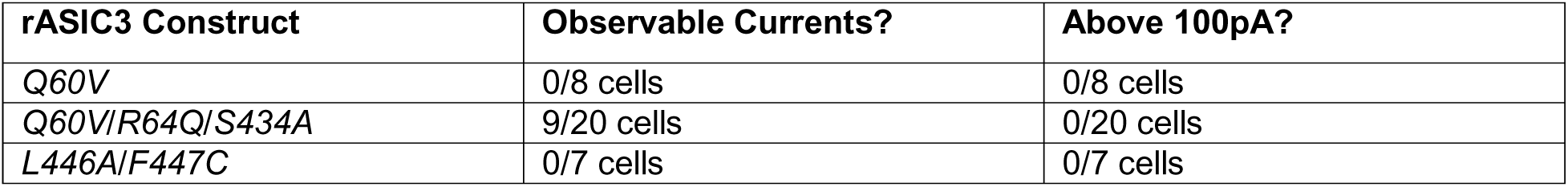
Non-functional/ poorly expressing mutants.

One residue in our simulation, hR68, is only present in the human orthologue. This arginine is not present in any of the ASIC isoforms or orthologues that have been confirmed to be regulated by single acyl chain lipid including human ASIC1a, rat ASIC1a, rat ASIC2a, and rat ASIC3 suggesting that it is not necessary for lipid potentiation of the channel^3,14,15^. To examine the other two arginine residues in rASIC3, rR64 (hR63) and rR66 (hR65), we mutated each to a glutamine to neutralize these sites. In addition, we made a double mutation of each residue to glutamine. All mutations produced functional channels that showed proton activated currents (Fig. 6A,C), although the channels that contain the rR64Q (hR63) mutation showed smaller current amplitudes. To assess the changes in channel desensitization rate, we measured the time it took for the current to decay from the peak of activation to 1/e for each mutant channel in the absence and presence of 10 or 20 µM DHA (Fig. 6B). When comparing the effects of DHA on each mutant channel, we found that the average decay rates for rR64Q/rR66Q double mutant channels did not change following the application of DHA at either concentration. When assessing the single-point mutant channels, R64Q was sufficient to eliminate the effect of DHA on channel desensitization at both concentrations while R66Q was not. However, rR66Q modestly reduced the effect of the lipid suggesting that both interactions with surface and buried sites might play some role in the effects of DHA on ASIC3 channel desensitization (Supplementary Table 2). In combination with our MD simulations, these findings indicate that the exposure of hR63 upon channel opening creates a more stable interaction site for DHA that is critical for the ability of the lipid to alter channel desensitization, while hR65 and hR68 form an adjacent electropositive surface that may facilitate DHA recruitment.

**Figure 6.**
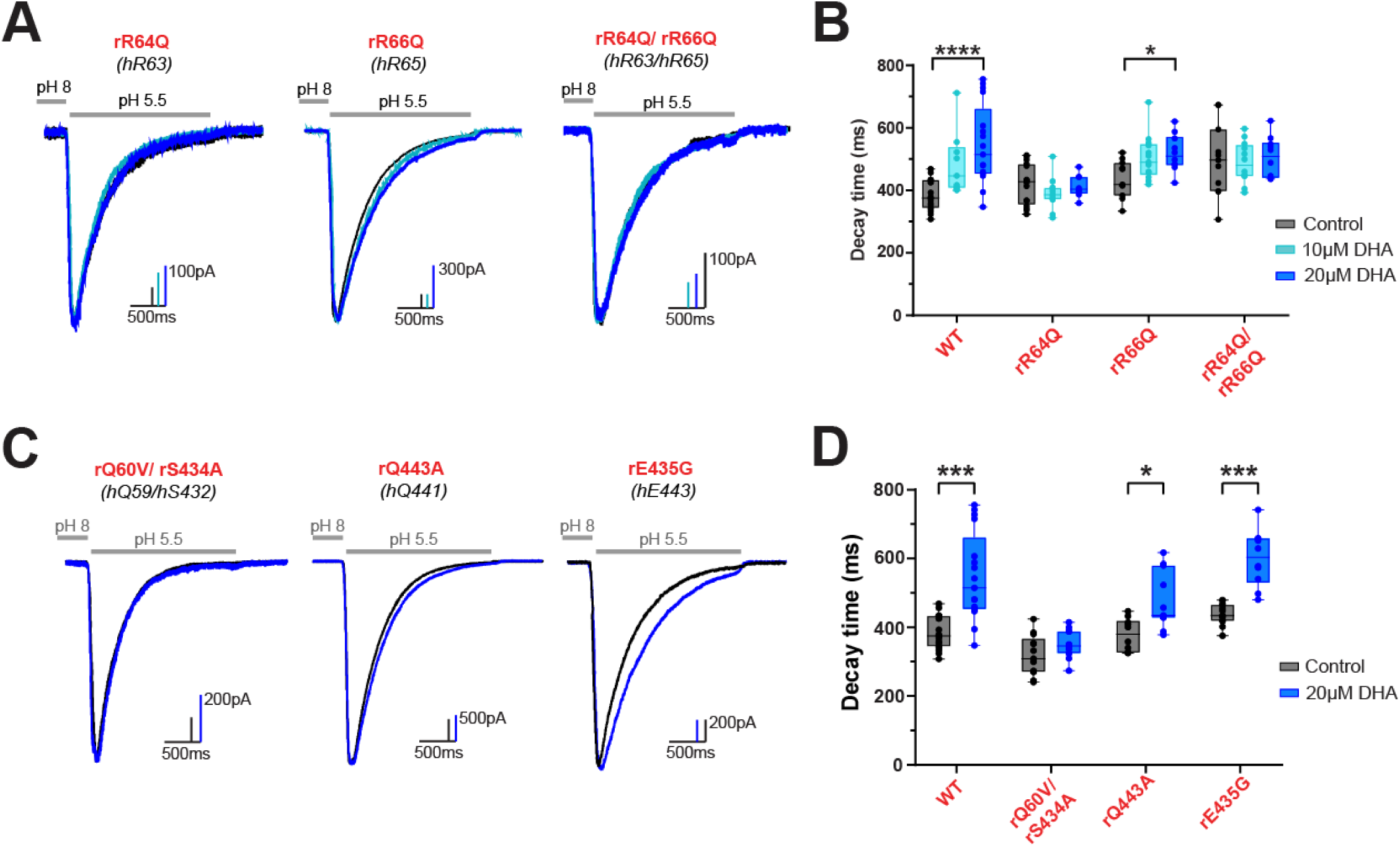
Mutating the PUFA binding site prevents slowing of the desensitization rate. **(A)** Representative whole-cell recordings showing the slowing of the decay rate in ASIC3 mutants ± 10 and 20 μM DHA. **(B)** Box plot showing the time point at which ASIC3 mutant currents decayed from their peak by 63% (*<u>R64Q:</u>* 0 μM = 418.9 ± 16.2 ms, *n = 16*; 10 μM = 389.8 ± 13.4 ms, *n = 13*; 20 μM = 413.63 ± 13.1 ms, *n = 8. <u>R66Q:</u>* 0 μM = 435.4 ± 17.8 ms, *n = 11*; 10 μM = 504.0 ± 19.7 ms, *n = 13*; 20 μM = 519.2 ± 18.7 ms, *n = 10. <u>R64Q/R66Q:</u>* 0 μM = 497.7 ± 35 ms, *n = 11*; 10 μM = 488.9 ± 19 ms, *n = 12*; 20μM = 505.4 ± 20.7 ms, *n = 10.*). **(C)** Representative whole-cell recordings showing the slowing of the desensitization rate of ASIC3 mutants ± 20µM DHA. **(D)** Box plot showing the time point at which ASIC3 mutant currents decayed from their peak by 63% (1/e) at 0 and 20 µM DHA (*<u>Q60V/S434A:</u>* 0 μM = 318 ± 16.1 ms, *n = 13*; 20 μM = 349.4 ± 11 ms, *n = 13. <u>Q443A:</u>* 0 μM = 377.7 ± 15.1 ms, *n = 10*; 20 μM = 477 ± 25.4 ms, *n = 11. <u>E435G:</u>* 0 μM = 438.4 ± 8.5 ms, *n = 13*; 20 μM = 601 ± 25.9 ms, *n = 10.*). All data given as mean ± SEM. C, One-way ANOVA post hoc Dunnett’s test. E, unpaired t-test with Welch’s correction. *, P < 0.05; **, P < 0.01; ***, P < 0.001; ****, P < 0.0001. ASIC3 WT data are replotted from Fig. 1 for comparison.

We continued to examine our putative interface by mutating additional sites that appeared to be critical for stabilizing the lipid at the more buried binding site. Here we only compared the desensitization rates from currents elicited at pH 5.5 in the absence and presence of 20 µM DHA. As a negative control for the model, we also chose to mutate a residue located near rR64 (hR63) in rat ASIC3 but showed little to no interaction with the head group of DHA throughout the simulations, rE435 (hE443). In agreement with the model’s prediction of a critical buried binding site, mutations to rQ60 (hQ59) and rS434 (hS432) together in rASIC3 significantly reduced the effect of DHA on the desensitization rate of the channel, only producing a slight change in decay rate by 30 ms (Fig. 6C, D; Supplementary Table 2). The individual mutants at these two positions resulted in small or absent currents preventing the determination of whether both residues are critical or just one of the two mutated here. rQ443 (hQ441) mutant channels made a very modest but not significant reduction in the effect of DHA Meanwhile, for the mutation of E435 to glycine DHA produced a 162 ms slowing in the decay rate, which is comparable to the 168ms slowing that is produced in WT channels.

Interestingly, the effect of DHA on the desensitization rate correlates well with the overall interaction time observed in our simulations: mutation of the residues with the longest overall interaction times (most contacts and longest durations) showed an either total elimination or near total elimination of the DHA-mediated regulation. Residues with more modest interactions, like rQ433 (hQ441) and rR66 (hR65) showed more modest impacts of mutations on the DHA effect. Taken together, these results suggest that this buried binding site, comprised of hR63, hQ59, hS432, and hQ441 is critical for the DHA-dependent slowing of channel desensitization.

### Identified DHA-ASIC3 interactions do not explain effects on pH-dependent channel activation

Single acyl chain lipids that regulate ASIC3 exert multiple effects on the channel. In addition to slowing desensitization, we have shown that many PUFAs shift the pH dependence of activation to more basic pHs. In principle, DHA could potentiate both effects through lipid interactions within the same region that we have now identified near the outer leaflet of the TMs. To test this, we examined the effect that 20 µM DHA has on the pH dependence of activation in our group of ASIC3 mutants. Like what we have shown for WT, we plotted pH activation curves using currents from ASIC3 mutants that were elicited by brief pulses of varying pH solutions in the absence and presence of DHA (Supplementary Figure 8) and determined the half activating pH by fitting the curves to the Hill equation (Fig. 7A-B).

**Figure 7.**
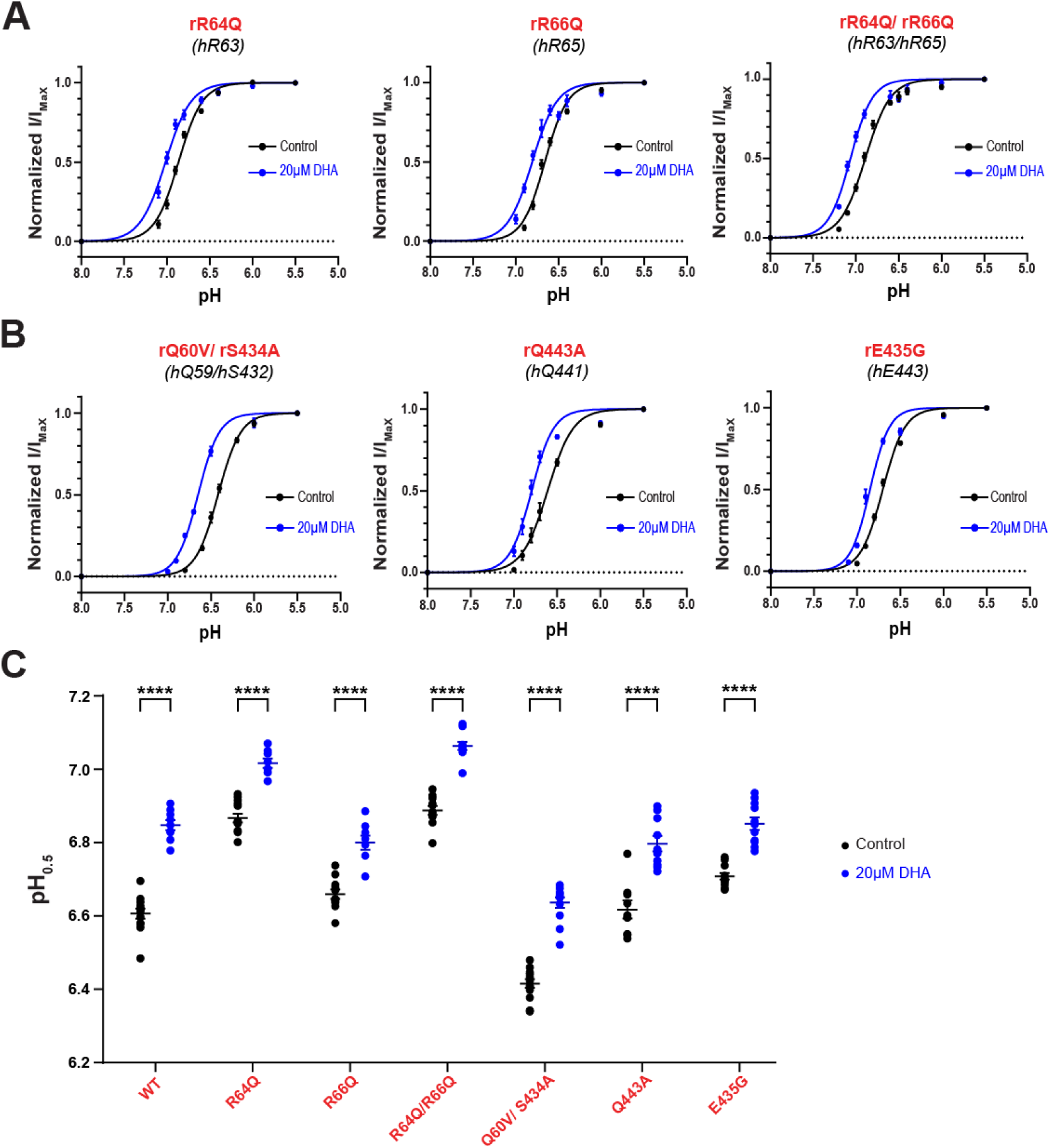
Mutations to TM residues are still impacted by the effect of DHA on pH-dependent channel activation. Fitted curves represent the pH dependence of channel activation of ASIC3 arginine mutants **(A)** and contact analysis mutants **(B)** ± 20 µM DHA, fitted using a Hill-type equation (see Equation 1 in Methods). **(C)** Average activation pH_0.5_ values for ASIC3 mutant channels in response to 0 and 20µM DHA (*<u>R64Q:</u>* 0 μM = 6.87 ± 0.012 pH, *n = 13*; 20 μM = 7.02 ± 0.012 pH, *n = 8; <u>R66Q:</u>* 0 μM = 6.66 ± 0.013 pH, *n = 11*; 20 μM = 6.80 ± 0.019 pH, *n = 8. <u>R64Q/R66Q:</u>* 0 μM = 6.89 ± 0.012 pH, *n = 11*; 20 μM = 7.06 ± 0.011 pH, *n = 11. <u>Q60V/S434A:</u>* 0 μM = 6.42 ± 0.012 pH, *n = 11*; 20 μM = 6.64 ± 0.014 pH, *n = 13. <u>Q443A:</u>* 0 μM = 6.62 ± 0.025 pH, *n = 9*; 20 μM = 6.80 ± 0.021 pH, *n = 10. <u>E435G:</u>* 0 μM = 6.71 ± 0.009 pH, *n = 12*; 20 μM = 6.85 ± 0.017 pH, *n = 11*). ASIC3 WT data are replotted from Fig. 1 for comparison. All data given as mean ± SEM. C, unpaired t-test with Welch’s correction. ****, P < 0.0001.

Based on our findings thus far, we hypothesized that mutant channels that had a robust effect on disrupting DHA-mediated slowing in channel desensitization should also disrupt DHA’s ability to shift the pH dependence of channel activation towards more basic pHs. However, surprisingly, all tested mutations still display significant shifts in the average pH_0.5_ values following application of 20 µM DHA (Fig. 7C). However, comparison of the magnitude of the shifts suggests there might be trend toward a modest reduction in the shift in proton sensitivity for rR64Q, rR66Q, and E435G (Supplementary Table 2). It’s possible that this gating effect requires a more complex set of interactions that if we could make multiple mutations, we would eliminate this effect. We attempted to make a triple-point mutation to Q60, R64, and S434; however, this proved to be detrimental to channel expression and function (Table 2). It is also possible that this network of interacting residues prevents simple mutants from eliminating interactions between DHA and ASIC3, and that the lipid alters channel function through a change in the local environment around the channel and not through a ligand-like interaction. Altogether though, this data suggests that binding to this region of charged and polar residues along the outer leaflet of the TMs largely does not eliminate the ability of DHA to produce alkaline shifts in the pH dependence of channel activation.

### NAAAs and LPCs may utilize a different binding mechanism to modulate ASIC3

We and others have previously shown that ASIC3 can be modulated by at least three classes of lipids: PUFAS, NAAAs, and LPCs. NAAAs, like AG and arachidonoyl serine (AS), have even more potent effects on slowing the rate of channel desensitization when compared to PUFAs while LPCs produce the largest effects out of any lipid tested on ASICs thus far^3,14^. All three classes of lipids produce a slowing in the desensitization rate and an alkaline shift in the pH dependence of activation. It remains unclear though if these lipids, despite their structural differences, produce their functional effects through similar interaction mechanisms.

To test this, we measured the effects of multiple lipids on R64Q mutant channels (Fig. 8A-B). We measured current upon solution switch from pH 8 to pH 5.5 and determined the time it takes the current to decay to 1/e of the peak value. Again, we preincubated our cells for at least 5 minutes with the indicated lipid prior to measuring currents. First, we tested three different PUFAs. AA and DHA are the most common PUFAs found within neuronal membranes, and both caused a significant slowing of ASIC3 WT desensitization (Fig. 8B-C, Supplementary Table 1). Surprisingly, eicosatrienoic acid (ETA) only showed a non-significant trend towards slowing desensitization rate despite previous work from our lab showing that it had the second largest effect on shifting ASIC3 proton sensitivity^3^. R64Q mutant channels showed no response to application of any of these three PUFAS at 20µM concentrations. We then tested two NAAAs (AG and AS), and both showed a robust slowing of channel desensitization in both WT and R64Q mutant channels (Fig.8B-D, Supplementary Table 1). Unexpectedly, R64Q channels showed an increased response to AS. Previous work identified two lysophosphatidylcholines, LPC(18:1) and LPC(16:0), as strong potentiators of ASIC3. Here we applied a mixture of LPCs which mainly contained the species LPC(16:0) to cells expressing either WT or R64Q mutant channels and found that both showed a dramatic slowing of channel desensitization. Like AS, LPC had an even larger effect on R64Q mutant channels compared to WT (Fig 8B-D, Supplementary Table 1). Because the head groups of NAAAs and LPCs tend to be larger and more electrically charged compared to that that seen in PUFAs (Fig. 8A), it’s possible that they utilize other interactions more within this region to stabilize themselves at the binding site. However, neutralizing other polar residues identified within this putative region did not prevent AG-mediated effects on channel desensitization rate or pH dependence for channel activation either although a trend towards a reduced effect can be seen for rQ60V/rS434A and rQ443A (Fig. 8E-F, Supplementary Figure 8; Supplementary Table 3, 4). Altogether, our data suggests that the state dependent interaction with an arginine in TM1 is required for PUFAs to slow the desensitization rate of the channel but is not sufficient to explain how other potentiating lipids cause the same general gating change. Moreover, the shift in the pH dependence of activation does not require interaction with that arginine either.

**Figure 8.**
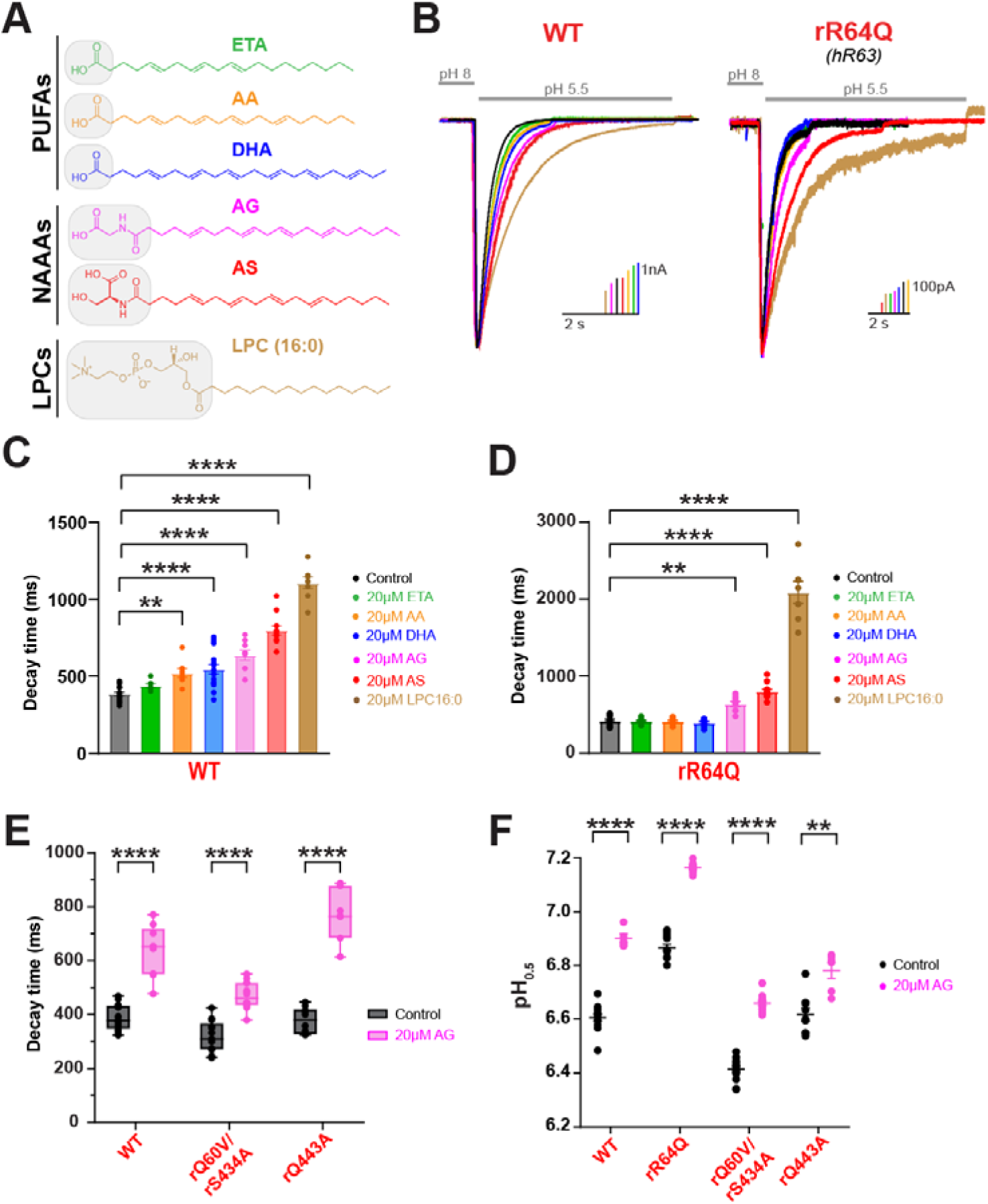
Mutations to TM residues disrupt PUFAs from slowing ASIC3 desensitization but not NAAAs of LPCs. **(A)** Chemical structures of PUFAs (ETA, AA, and DHA), NAAAs (AG and AS), and LPCs (LPC 16:0, which contain a 16-carbon chain with 0 cis double bonds) which were used in the following experiments. **(B)** Representative whole-cell recordings showing the slowing of the desensitization rate of ASIC3 WT (left) or R64Q (right) ± 20 µM lipid. Currents were elicited by performing fast-switch perfusion changes between pH 8 and pH 5.5 solutions. **(C)** Bar graph showing the average time point at which ASIC3 WT currents decayed from their peak by 63% (1/e) at 0 and 20 µM lipid. ASIC3 WT control and DHA data are replotted from Fig. 1 for comparison. **(D)** Bar graph showing the average time point at which ASIC3 R64Q currents decayed from their peak by 63% (1/e) at 0 and 20 µM lipid. ASIC3 R64Q control and DHA data are replotted from Fig. 4 for comparison. **(E)** Box plot showing the average time point at which ASIC3 WT and contact analysis mutant currents decayed from their peak by 63% (1/e) ± 20 µM AG application. ASIC3 WT control and AG data are replotted from Fig. 7C for comparison. **(F)** Average activation pH_0.5_ values for ASIC3 WT and mutant channels in response to 0 and 20 µM AG. All data given as mean ± SEM. C-D, One-way ANOVA post hoc Dunnett’s test; E-F, unpaired t-test with Welch’s correction. *, P < 0.05; **, P < 0.01; ****, P < 0.0001 (see Methods and Supplementary Table 1, 3, and 4 for details).

## Discussion

To date, there have been several studies showing the physiological consequences of lipid potentiation on ASIC currents^14–16,22,34^; by shifting the pH dependence for channel activation towards more physiological concentrations, neurons become more sensitized to firing action potentials following mild acidosis and in some cases in the absence of acidosis^14,16^. In addition to changes in proton sensitivity, there are also significant changes to channel desensitization rate. Here we examined the interaction of DHA with ASIC3 using MD simulations and then determined the functional relevance of those interactions using site-directed mutagenesis and patch-clamp electrophysiology. We found that PUFAs interact with the channel in the resting state at a positively charged patch on the lipid-facing side of TM1. Upon channel opening, a more buried binding site is exposed and interactions at this site were critical for the slowing of channel desensitization by DHA. In addition to this first ever description of a functional binding site for lipids on ASICs, our work made three surprising discoveries. First, mutation of the more exterior site residues, like hR65, had modest impacts on lipid-mediated slowing of channel desensitization and proton sensitivity. Second, interactions the eliminated the lipid effect on channel desensitization had only mild effects on the pH dependence of channel activation. Finally, mutation of the interaction site did not impact the ability of other classes of lipids known to potentiate ASICs including NAAAs and LPC and in some cases paradoxically increased the effect of the lipid.

Our description of the binding interface here is largely consistent with growing evidence in the field. Mutations in and around the extracellular surface of TM1 were previously shown to reduce the effect of PUFAs on ASIC3^3^. This region included the arginine residue hR63 (R64 in rat ASIC3) that we identify in this paper. Interestingly, hR63 was not predicted to be involved in arachidonic acid binding *in-silico*, creating some discrepancies between computational and experimental data^29^. This previous computational study suggested that hR65 and hR68 were the critical binding site for arachidonic acid on ASIC3. Like these previous simulations, we found that these same positively charged arginine residues which comprise the more surface lipid binding site and face towards the membrane environment attract the negatively charged carboxyl head group of DHA molecules in both the resting and open state. However, hR68 is not present in the rat orthologue and mutation of rR66, the other exterior arginine, had only a small effect on the lipid modulation of the channel. Despite this, the more exterior site may result in an increased occupancy of DHA molecules near the channel, suggesting that the local lipid environment surrounding ASIC3 channels becomes enriched with DHA compared to the bulk membrane. Unlike previous work, we found that when the channel transitions into an open state, DHA makes much longer-lived interactions with an arginine that is only exposed in the open state (hR63). Helping to stabilize this interaction, the head group also makes contacts with nearby polar residues including hQ59 and hS432. We hypothesize, based on these results, that this state-dependent interaction is the mechanism by which DHA stabilizes the open state of ASIC3 by slowing the rate at which channels transition from a conducting state into a non-conducting state. In addition, although ASIC3 exhibits C3 symmetry, DHA occupancy in our simulations is not perfectly symmetric across the three subunits. This likely reflects the stochastic nature of lipid diffusion and binding, as well as the finite simulation timescale, which may limit equal sampling of equivalent sites. Importantly, we observe similar interaction patterns across subunits, suggesting that the identified binding region represents a general feature of the channel rather than a subunit-specific effect. While we cannot say for sure why our simulations found this stable interaction but previous ones did not, it may arise from the fact that our all-atoms MD simulations were simulated for 5µs, which provides a 10-fold longer sampling rate than what has been shown in previous work^29^. While the interaction with hR63 is more stable, it also occurs less frequently, leaving open the possibility that this interaction is less likely to be detected during shorter simulations. A limitation of this study is that the simulations are based on homology models of ASIC3, as no experimental structures are currently available. While the models were constructed using the best available templates and are consistent with independent structural predictions, some uncertainty remains, particularly in the positioning of side chains at the lipid-facing interface. Notably, hR63 is conserved across species, providing additional confidence in conclusions involving this residue.

While this study has focused primarily on PUFAs, there are other classes of single acyl chain lipids that we and others have shown to produce a slowing effect on the desensitization rate and an alkaline shift in the pH dependence of activation^3,14^. Given the impact that each of these classes of lipid has on channel function is qualitatively similar, it would be easy to hypothesize that these lipids share a common binding surface. However, we found that neutralizing the critical arginine, R64 in rat ASIC3, was only capable of disrupting PUFAs from slowing the rate of channel desensitization but was not sufficient to disrupt this effect produced by NAAAs or LPCs. Moreover, neutralizing other polar residues identified within this putative region did not prevent AG-mediated effects on channel desensitization rate or pH dependence for channel activation. Surprisingly, the rR64Q mutant channels showed an even larger effect on the slowing of desensitization rate following the application of AS and LPC. There are several potential explanations for this result. First, the idea that there is a common binding region for these lipids could be wrong. There may be novel binding sites that will be uncovered in future simulations of AG and LPC binding. However, we hypothesize that, given the common functional impact, that the lipids may use a shared binding surface on the channel but that the precise residues involved vary due to the unique structures of the head groups. These lipids may be attracted to this region of the channel through the positively charged arginine residues on the top of TM1 but then make their own set of interactions with residues in this region based on head group structure. The more buried arginine residue may be important for each of these interactions, but the larger and more complex head groups of NAAAs and LPCs may prevent a single point mutation from eliminating lipid mediated regulation. NAAAs have head group structures that contain the same carboxyl group as PUFAs but additional reactive groups. Arachidonoyl Serine, for instance, has an additional amino group and hydroxyl group that could increase its interaction surface which could prevent a single point mutation from being able to eliminate binding of these lipids to the channel. LPC is zwitterionic with both a phosphate and choline as part of its head group creating a much larger surface for potential interactions. In addition, AS and AG head groups are more electronegative than PUFA headgroups and LPC is thought to create a negative charge at the membrane/aqueous interface consistent with the notion that these lipids might be attracted to a positive region at the top of TM1^35,36^.

While the binding interface identified here appears to be necessary for the ability of the lipid to slow channel desensitization, it does not fully explain the change in proton sensitivity that occurs additionally. Mutations to this binding site were still able to produce shifts in the pH dependence in the basic direction following DHA application, suggesting that another mechanism may be involved. One possibility is the presence of a second binding site. Although we did not identify another external binding site within our simulations that were more stable than R63, it’s possible that PUFAs modulate different gating effects independently through multiple interaction sites. Work examining PUFA regulation of Kv7.1 channels has described a similar mechanism where differing gating effects are controlled through interactions with different sites on the channel^30^. Alternatively, this effect could be driven through a more complex set of interactions between PUFAs and ASIC3 at this binding interface, including those that are occurring along the tail group of the lipid. We previously found that PUFA potentiation of ASIC3 was largely driven by the electrostatic potential of the head group, however the properties of the tail group also had some impact on the potency of the lipid^3^. Within our simulations and others^29^, the PUFA carbon tails appear to position themselves along the TMDs. Protein-lipid interactions at these sites may serve to provide structural support for PUFAs to remain near the channel or to alter gating. We attempted several mutations at sites that our simulations showed interactions between the tail groups and the channel but mutating those residues resulted in nonfunctional channels.

It is also possible that the accumulation of DHA near the channel alters channel function through solvent-like effects on channel function. Solvent-like has been defined as non-ligand-like effects that occur through changes in local properties like bilayer fluidity or thickness. Many ion channels including Piezo channels are notably sensitive to these sort of membrane physical properties^37^. Our data does not suggest which solvent-like effect may be impacting the apparent sensitivity of the channel to protons. However, we have previously showed that the electronegativity of the lipid head group is critical for determining the magnitude of the effect of the lipid on ASIC3 function^3^. This increased electronegativity around the lipid-aqueous interface could have significant effects on the local pH near the channel^35,36^; negatively charged lipids surrounding the channel may accumulate more protons, which could result in a decrease in the local pH immediately surrounding the channel compared to the bulk pH. This would ultimately manifest as a shift in the pH dependence of channel activation in the basic direction. This also may explain the inability of our point mutations to eliminate the impact of DHA on the pH dependence of activation. Future work using fluorescence approaches to examine this question of changes in juxtamembranous pH could help answer this question^38^. In addition, the use of lipids to modulate membrane physical properties will also be essential to examine the question of whether ASIC3 is impacted by both ligand-like and solvent-like effects.

Our mutation of R64Q in this study is not completely consistent with our previous result that did show a significant reduction in the impact DHA has on the proton sensitivity of the mutant channel. However, the data here shows a trend towards a reduction in the effect that DHA has on the pH dependence of activation, and our previous work shows a significant but only partial reduction in the effect. We think that, likely, mutation of the more stable interaction sites on the channel could reduce the accumulation of these lipids around the channel and thus reduce the possible solvent-like effect on channel function.

Increase of single acyl chain lipids like PUFAs and LPCs in the plasma membrane occurs in joint inflammation and leads to ASIC3-mediated pain^14^. Additionally, patients with rheumatic diseases that experience chronic pain and have in increased level of LPC in their synovial fluid are thought to feel pain through a mechanism involving ASIC3 potentiation via lipid^34^. Our description of a binding region for single acyl chain lipids on ASIC3 opens the door to searching for molecules that would inhibit binding of these lipids to this region of the channel and potentially reduce ASIC-mediated pain during joint inflammation. Our work here describes a relatively small binding site made up of a series of somewhat diffuse interaction sites for PUFAs on ASIC3 consistent with the idea that these lipids are likely acting through a combination of ligand-like and solvent-like effects. A competitor that blocked this region could eliminate all the potentiating effects of these lipids on ASIC3. However, more work is needed to understand of the other classes of potentiating lipids are acting through this same binding surface.

## Methods

### Materials and mutagenesis

Docosahexaenoic acid (DHA; Cat. # 90310), N-Arachidonyl Glycine (AG; Cat. # 90051), Arachidonic Acid (AA; Cat. # 90010), 5(Z),8(Z),11(Z)-Eicosatrienoic Acid (ETA; Cat. # 90190), N-Arachidonoyl-L-Serine (AS; Cat. # 10005455) and Lysophosphatidylcholine (LPC; Cat. # 24331) were purchased from Cayman Chemical for use throughout all electrophysiology experiments. Rat ASIC3 plasmid was gifted by David Julius (University of California, San Francisco, San Francisco, CA) and subcloned into a pcDNA3.1 vector. The fluorescent tag mCerulean3 was attached to the C-terminus of the channel using a short proline rich linker to monitor its expression levels, which has been previously reported to have minimal effects on channel gating^39^. All mutant rat ASIC3 channels used in this study were made using site-directed mutagenesis either in-house (KOD Hot Start Master Mix; Cat. #71842) or through Biozilla services. Successful mutagenesis was verified through whole plasmid sequencing services through Plasmidasaurus or sanger sequencing services through ACTG. Mutant plasmids were then maxi-prepped using endotoxin-free methods either in-house (Qiagen; Cat. # 12362) or through plasmid DNA purification services through Biozilla.

### Cell culture and Transfection

CHO-K1 cells (ATCC) were cultured using Ham’s F12 medium supplemented with 10% FBS and were incubated at 37°C in 5% CO_2_. Cells grew to 70-80% confluency before being transfected with rat ASIC3 WT or mutant plasmid DNA (1.5-3µg). This was done either by electroporation with a Lonza 4D Nucleofector unit (Cat. # V4XC-1032) or by chemical transfection using TransIT transfection reagents (Cat. # MIR 2304) according to each manufacturer’s protocols. Following transfection, cells were plated on 12-mm glass coverslips coated in poly-L-lysine and incubated at 30°C in 5% CO_2_.

### Preparation and application of lipids

All electrophysiological experiments which were in the presence of lipids were performed under identical conditions as control experiments except the resting pH 8 solutions contained the indicated concentration of lipid. All lipid stock solutions were made up in ethanol except in the case of LPC which was made up in a 2:1 ratio of chloroform: methanol (LPC) following the manufacturer’s recommendations. Ethanol or chloroform: methanol solvent in final solution was typically 0.01% and never exceeded 0.1%. Solution pH was measured prior to and after addition of PUFAs to solution to ensure no pH change occurred. For experiments that included lipid, cells were exposed to a holding pH 8 solution containing the indicated concentration of lipid for 5 minutes prior to eliciting pH activations unless stated otherwise to ensure any possible effects that were present under each condition would be saturated.

### Electrophysiological recordings

As previously described^3,40^, all experiments were performed 16–30 h after transfection. To assess peak current magnitudes, whole-cell patch-clamp configuration was used. Borosilicate glass pipettes (Harvard Apparatus) were pulled to a resistance of 2–4 MΩ (P-1000; Sutter Instrument) for whole-cell experiments. Glass pipettes were filled with an internal solution containing (in mM) 20 EGTA, 10 HEPES, 50 CsCl, 10 NaCl, and 60 CsF, pH 7.2. Extracellular solution contained (in mM) 110 NaCl, 5 KCl, 40 NMDG, 10 MES, 10 HEPES, 5 glucose, 10 Trizma base, 2 CaCl2, and 1 MgCl2, and pH was adjusted with HCl or NaOH as needed. An Axopatch 200B amplifier and pCLAMP 10.6 (Axon Instruments) were used to record whole-cell rASIC3 WT and mutant currents. All recordings were performed at a holding potential of −80 mV with a 5-kHz low-pass filter and sampling at 10 kHz. Solution changes were performed using a rapid perfusion system (SF-77B Fast-Step perfusion, Warner Instruments). The fluorescence of each recorded cell was visualized on an Olympus IX73 microscope with a CoolLED pE-4000 illumination system.

#### Desensitization rate

To measure desensitization rates in the absence and presence of DHA and other PUFAs, cells were exposed to a holding pH of 8 for 8 s with or without lipid followed by a 2-s application of pH 5.5. For experiments on WT channels including AS and LPC applications, cells were exposed to a holding pH of 8 for 14 s followed by a 6-s application of pH 5.5. For experiments on R64Q channels including AS application, cells were exposed to the same conditions as WT channels while those including LPC were exposed to a holding pH of 8 for 20 s followed by 8-s applications of pH 5.5 to ensure the current decay was mostly saturated. Exposure to pH 8 solution between pH 5.5-activating pulses was also increased to ensure full recovery of the channels expressed on the cell membrane between each sweep. Rates were determined by taking the time point at which 1/e (63%) of the peak of pH 5.5 current had decayed. Each individual experiment was measured across multiple sweeps that were then averaged to obtain an average decay time for that experiment.

#### Shifts in pH-dependence of activation

To determine the pH dependence of channel activation, a series of sweeps were measured from a holding pH of 8 for 8 s, followed by 2-s activation pulses at indicated pHs starting from a maximally activating pH of 5.5 down to a minimally activating pH which was dependent on construct and lipid condition. Each activation pH was measured for two sweeps which were averaged together and then were normalized to the responses elicited at pH 5.5.

#### Wash-on rate

To measure the increase in current amplitude over time, cells in the absence of PUFAs were exposed to a holding pH of 8 for 4 seconds followed by a 1-s minimally activating pulse of pH 6.8. Cells were only exposed to lipid conditions at the initiation of each experiment, except in the case of experiments where internal application of lipid was achieved. In this case, cells were exposed to internal application of lipid for no more than 30 seconds prior to the start of each experiment to ensure that seals remained stable following break in. Data was normalized by comparing each of the responses within each experiment to the peak current within the first sweep.

### Data and statistical analysis

PUFA structure schematics provided within the figures were created using MarvinSketch chemical editing software (ChemAxon). ClustalW multiple sequence alignment was performed using Jalview 2.11.5.0 software and manually edited in Adobe Illustrator (2025). Sequences were retrieved from Uniprot using the following Uniprot IDs: Q9UHC3, human ASIC3; O35240, rat ASIC3; Q62962, rat ASIC2; P55926, rat ASIC1; P78348, human ASIC1. Whole-cell patch clamp current recordings were analyzed using Clampfit 10.6 (Axon Instruments). All statistical data are plotted as mean ± SEM.

For pH dependence, reported pH_0.5_ values represent the mean values for each individual experiment as determined by fitting to a Hill-type equation in GraphPad:

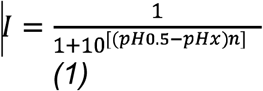

where *n* is the Hill number and pH_0.5_ equals the half-maximal activating pH. Plotted fits were restrained to a maximum value of 1 and a minimum value of 0.

Statistical analyses were performed using GraphPad Prism 10 software. P values are reported in tables as calculated except for p-values < 0.0001, where they are reported as such. Figure legends indicate which statistical test was performed for each data set. For figures, statistical significance is indicated by the following scale: *, P < 0.05; **, P < 0.01; ***, P < 0.001; ****, P < 0.0001. All p-values reported account for multiple comparisons using appropriate tests. For all statistical testing requiring multiple comparisons within each construct, a One-way ANOVA was performed followed by post hoc testing. P values reported in Supplementary Tables 1-3 represent Dunnett’s multiple comparisons test performed on all lipid data sets in the table compared with the control ASIC3 dataset. Unpaired student’s t tests were performed where indicated for statistical testing between two sets of data within each construct (e.g., Supplementary Table 4). All t tests were two-tailed, and unpaired t tests assumed unequal variance.

### Homology Modeling and Molecular Docking

Since no experimentally resolved structures for human ASIC3 are available in the Protein Data Bank (PDB), homology modeling was used to construct its structure in the open state. The chicken ASIC1 structure in the open state (PDB ID: 4NTY) and the human ASIC1 structure in the resting state (PDB ID: 7CFS) were employed as templates for modeling. The protein sequence of hASIC3 was retrieved from the UniProt database (UniProt ID: Q9UHC3) and submitted to the Swiss-Model web server (version 1.2). Swiss-Model generated 12 homology models for the open and resting state of hASIC3. Model selection was based on the Global Model Quality Estimate (GMQE) score^41^, which provides a quality measure for predicted structures. The model with the highest GMQE score was selected to represent both the open and resting states of hASIC3. This structure was then used for molecular docking and for building simulation systems for molecular dynamics (MD) studies.

Molecular docking was performed in the Schrödinger suite^42^. Using the Protein Preparation Wizard, the protein structure was pre-processed by assigning bond orders, adding hydrogens, adjusting protonation states at pH 7.4, creating disulfide bonds, and removing water molecules. Hydrogen-bond optimization and restrained minimization were subsequently performed. The 3D structure of docosahexaenoic acid (DHA) was obtained from PubChem. LigPrep^42^ was used to generate possible ligand protonation states between pH 6.0 and 8.0 and to assign chirality. R63 was selected as the receptor grid center based on our first simulation in this study and previous studies identifying R63 as a critical residue for PUFA potentiation in rat ASIC3^3,29,40^. For the ligand midpoint box, dimensions were set to 30 × 30 × 30 Å. Docking was carried out using the Glide XP (extra precision) mode in Maestro^42^. The docked complexes were analyzed in the Pose Viewer, and the best-scoring poses (based on G-scores) were selected for MD simulations. Because ASIC3 is a homotrimer with three equivalent lipid-binding sites, three DHA molecules were docked to the R63 site of the open-state ASIC3 model and subsequently subjected to MD simulations to investigate binding interactions.

### Molecular Dynamics Simulation System Settings

All simulation systems were constructed using the CHARMM-GUI Membrane Builder^43,44^. Disulfide bonds were added between cysteine residue pairs (C92–C186, C164–C171, C282–C370, C315– C366, C319–C364, C328–C350, and C330–C342) to preserve the structural integrity and stability of the protein. Protonation states of ionizable residues were determined by pKa predictions from PROPKA3 at pH 7.4^45^, and residues were assigned to their default states accordingly. The protein was oriented along the bilayer normal (z-axis) using PPM 2.0 to ensure accurate alignment and helix tilt within the membrane^46^. The protein was then embedded into the lipid bilayer (Table 1), and each system was solvated in a 0.15 M NaCl aqueous solution using CHARMM-GUI protocols^43^. Trajectories 1 and 2, corresponding to the resting- and open-state hASIC3 models, were simulated in a POPC bilayer with a POPC:DHA ratio of 10:1, where DHA molecules were randomly distributed across both leaflets. Although this enrichment exceeds physiological concentrations, it enabled systematic sampling of DHA–protein contacts and reproducible quantification of lipid occupancy and interaction lifetimes without compromising bilayer stability. The simulation boxes for Trajectories 1 and 2 measured 103 × 103 × 142 Å³ and 105 × 105 × 138 Å³, containing approximately 200,000 and 182,000 atoms, respectively. For Trajectory 3, a 3-DHA–docked open-state hASIC3 structure was embedded in a pure POPC bilayer under identical solvation and ionic conditions, with a box size of 105 × 105 × 135 Å³ and a total of ∼175,000 atoms, allowing direct comparison of DHA binding events between spontaneous and pre-bound systems.

### All-atom MD Simulation protocol

All-atom MD simulations were performed using either NAMD 2.14^47^ or the specialized computational platform Anton2^48^. The CHARMM36m force field was used for proteins, lipids, and ions within the study^49–53^. The explicit representation of water molecules was achieved through the implementation of the TIP3P model^54^. The simulations were executed under NPT (constant number of particle N, pressure P, and temperature T) conditions at a temperature of 310 K and a pressure of 1 atm, utilizing periodic boundary conditions. Electrostatic interactions were handled using the particle mesh Ewald method with a 12 Å cutoff for real-space interactions^55^. Hydrogen atom bond distances were constrained through the application of the SHAKE algorithm^56^. Hydrogen mass repartitioning (HMR) allows us to re-distribute the mass of heavy atoms to the bonded hydrogens^57^. Hence a time step of 4 fs was selected for the simulations ran on NAMD. Prior to the production runs, minimization and equilibration were performed in NAMD 2.14 with harmonic positional restraints applied in two sequential stages. In the first stage, restraints were imposed on all protein heavy atoms, allowing solvent and lipids to relax around a fixed protein framework. In the second stage, the restraints were reduced to include only protein Cα atoms and ligand heavy atoms, permitting sidechain and lipid rearrangements while maintaining the overall backbone geometry. Each simulation had 5,000 steps of energy minimization followed by two-stage NPT equilibration for a combined duration of 2 ns. The equilibrated systems were simulated for 5 microseconds for the DHA partitioning simulations in both the open and resting states, while simulations with three DHA molecules docked were performed for 500 nanoseconds. In MD simulations conducted with Desmond on Anton2, temperature was controlled using a Nose–Hoover thermostat and pressure was controlled using the MTK barostat at 1 atm. The calculation of long-range electrostatic interactions was facilitated by the k-space Gaussian split Ewald method^58^, utilizing a 64 × 64 × 64 grid. Time-step for simulations run on Anton2 is 2.5 fs. MD trajectories were then visualized and analyzed using VMD^59^, in-house Tcl, and Python scripts.

#### Occupancy map of DHA

To quantify and visualize the preferred spatial distribution of DHA molecules surrounding the hASIC3 channel, time-averaged occupancy maps were generated using the VolMap plug-in in VMD 1.9.4^59^. Occupancy maps were computed for Trajectory 1 (resting state) and Trajectory 2 (open state) to capture state-dependent differences in DHA localization. The analysis focused on DHA carboxylate carbon atoms, which mediates electrostatic and hydrogen-bonding interactions with the protein surface. A 1 Å grid resolution was used, and the resulting volumetric density was normalized to yield an occupancy map, reflecting the frequency with which DHA head groups occupied each spatial grid element during the simulation. The maps were visualized at 0.3, corresponding to regions where DHA head-group atoms were present for at least 30 % of the total trajectory, revealing persistent DHA-binding zones near the arginine-rich region of hASIC3.

#### Binding lifetime analysis

Binding events were identified by computing, for each DHA molecule independently, the per-frame distance between its carboxylate carbon and all heavy atoms of the target protein residue, taking the minimum distance at each frame. To reduce high-frequency noise, a centered rolling average of 50 frames (∼50 ns) was applied to each distance trace before contact detection. A binding event was defined as a continuous interval during which this smoothed distance remained within 4.3 Å of the residue sidechain. Brief interruptions of fewer than 50 consecutive frames were merged into the same event to avoid artificial fragmentation due to transient fluctuations. This analysis was performed for residues Y58, Q59, R63, R65, R68, S432, and Q441 across all three subunits (chains A, B, C) in both the open and resting states. Event durations are reported in nanoseconds (1 ns per frame). For each residue–chain–state combination, the distribution of individual event durations is shown as a box-and-whisker plot (box: interquartile range; line: median; whiskers: 1.5× IQR); the longest individual binding event per group is highlighted with a star (★). Individual data points are shown as jittered dots overlaid on each box.

#### Contact analysis

Contact analysis was carried out using the inter-sel contacts function within the Timeline plug-in of VMD 1.9.4^59^. This analysis identified all residues of hASIC3 that formed close interactions with carboxylate carbon of DHA molecules throughout trajectory 3. A cutoff distance of 4 Å between heavy atoms was applied to define a contact, ensuring that both transient and stable interactions were captured. The contact number was calculated as the total number of atom pairs between DHA and each protein residue at a given time frame, providing a quantitative measure of lipid–protein interactions across the trajectory. This approach enabled the identification of residues most frequently involved in DHA binding.

## Acknowledgements

Research reported in this publication was supported by the National Institute of General Medical Sciences R35GM137912 (to JRB) and R35GM160133 (to JL). Additional support was provided by the National Science Foundation CAREER grant under award number 2439983. The content is solely the responsibility of the authors and does not necessarily represent the official views of the National Institutes of Health or National Science Foundation. Computer resources came from a Maximize ACCESS allocation through project BIO210015, an allocation (MCB200085P) on Anton2/Anton3 at the Pittsburgh Supercomputing Center, provided by the National Center for Multiscale Modeling of Biological Systems through National Institutes of Health grant P41GM103712-1, and from a loan from D. E. Shaw Research. The authors also thank the Computational Chemistry and Bioinformatics Research Core within the University of Mississippi’s Glycoscience Center of Research Excellence (NIH Project Number 5P20GM130460-04) for use of their computers and assistance with software installation.

## Contributions

RR performed the electrophysiology experiments and RB performed the MD simulations. All authors contributed to experimental design, analysis and project conceptualization. All authors contributed to data interpretation and the writing of the manuscript. JB and JL supervised the project and oversaw the acquisition of funding.

## Data Availability

The numerical source data underlying the simulation graphs and charts, as well as additional simulation data supporting the findings of this study, will be available on Zenodo (link will be updated after uploading). Due to their large file sizes, complete uncompressed molecular dynamics trajectories are available from the corresponding author upon reasonable request. Raw traces of functional recordings are available from corresponding author upon request. Representative raw traces appear either in the appropriate figure or in the supplementary data. Complete raw data sets can be requested as well but are not provided here due to their large and complex nature. The source data underlying the electrophysiology data is included here as a source data file.

## Supplementary Information

**Supplementary Figure 1.**
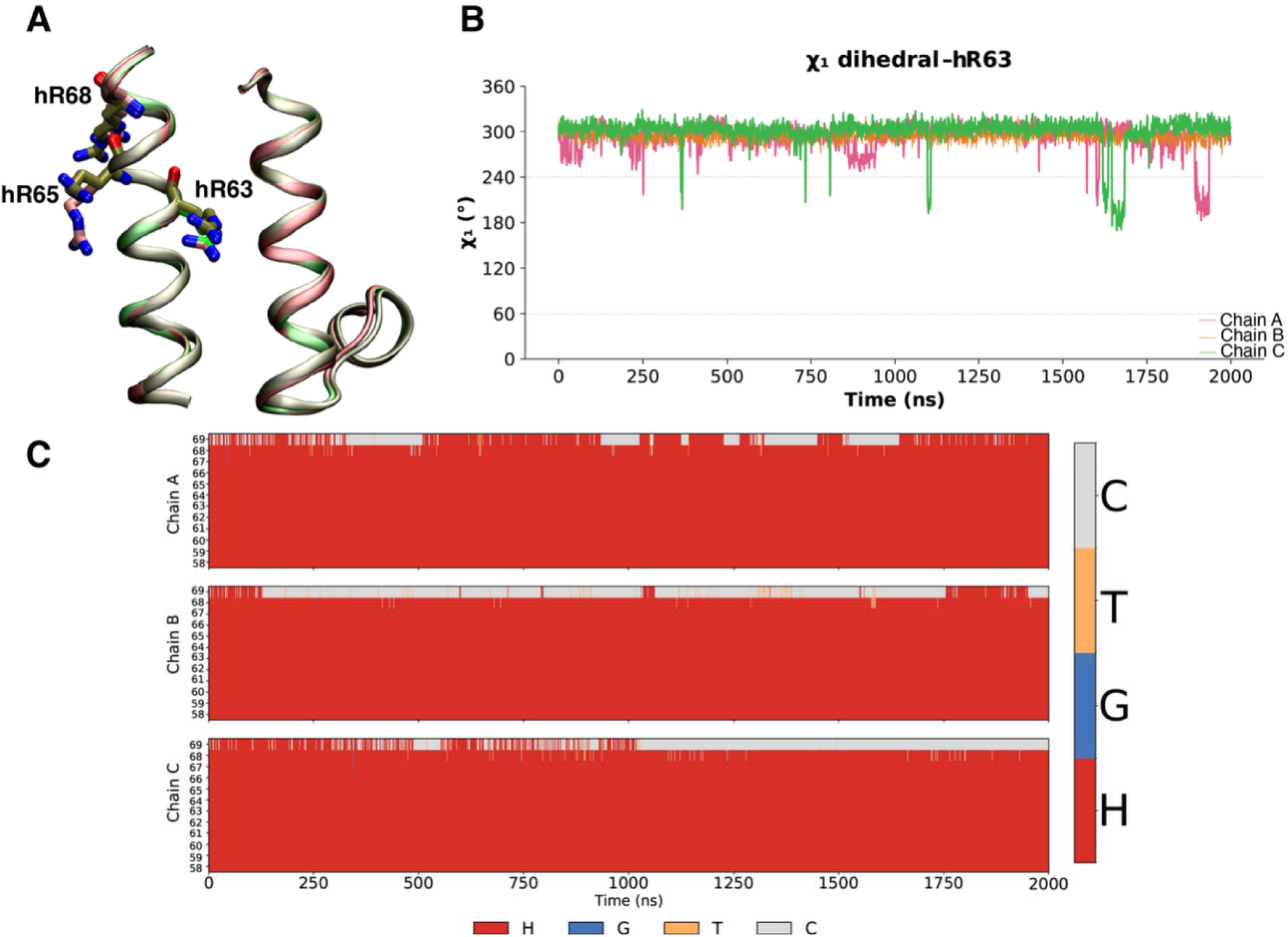
Structural alignment and conformational stability of key arginine residues in ASIC3. **(A)** Structural alignment of the transmembrane region showing key residues hR63 (rR64 and cR65), hR65 (rR66 and cR67), and hR68 (not an R in other isoforms). The **AlphaFold model (AF-Q9UHC3-4)** is shown in tan, the **homology model** in pink, and the **chicken ASIC1 open-state X-ray structure (PDB ID: 4NTY)** in green during Trajectory 4 (a ligand-free, apo production simulation of the open-state hASIC3 model; see Table 1), illustrating overall agreement in backbone architecture and side-chain positioning. **(B)** Time evolution of the χ₁ dihedral angle for hR63 across the three subunits (Chain A, pink; Chain B, orange; Chain C, green), showing a predominantly stable conformational state over the simulation with occasional transitions. **(C)** Secondary structure evolution of the transmembrane region over the course of the simulation for each subunit (Chains A–C), demonstrating overall structural stability. Secondary structure elements are colored as follows: helix (red), coil (gray), turn (orange), and 3₁₀-helix (blue).

**Supplementary Figure 2:**
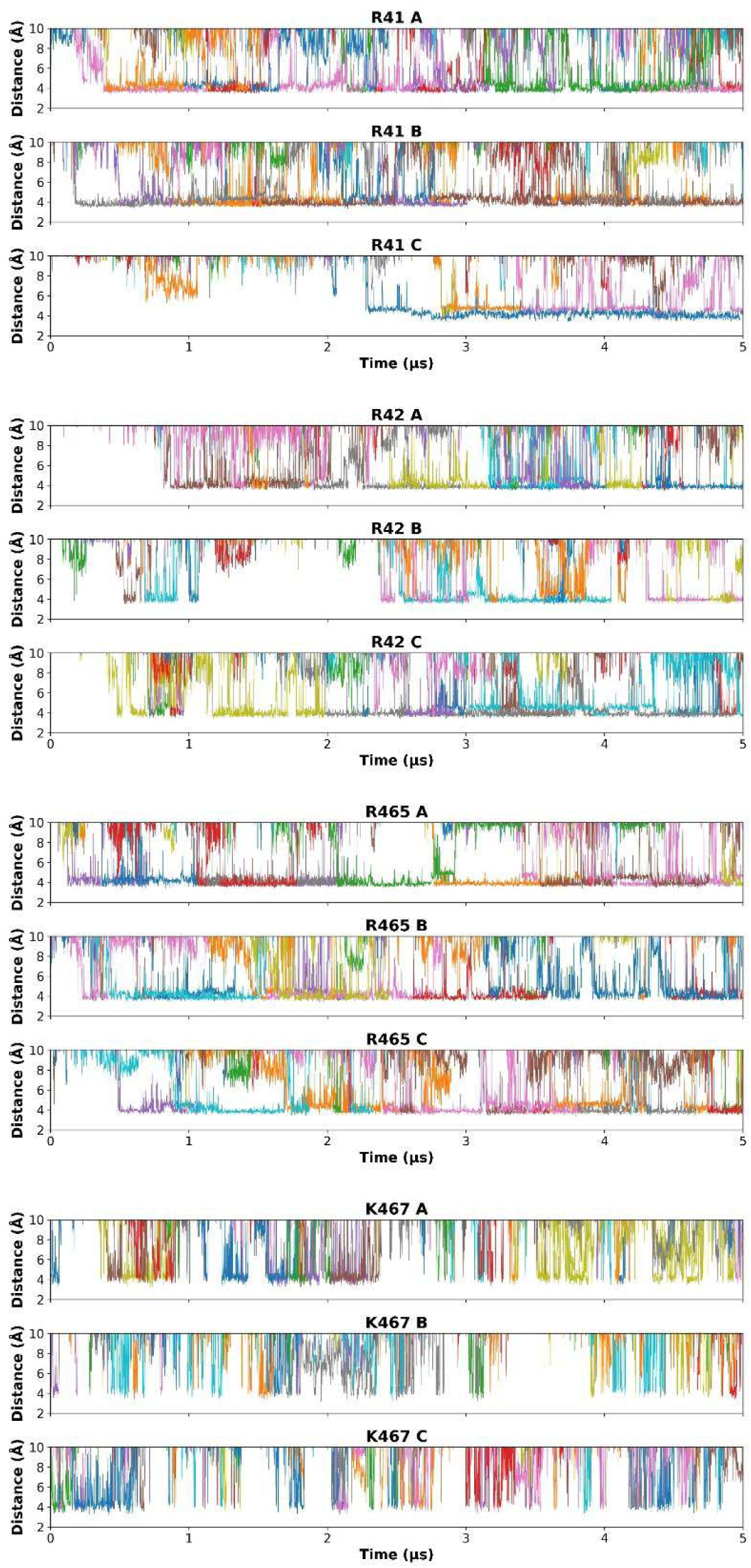
The spontaneous DHA binding to residues near inner leaflet in resting-state (Trajectory 1, the 5 µs simulation of the resting-state hASIC3 model with DHA randomly partitioned into the surrounding POPC bilayer; see Table 1): Distance plots showing the proximity between the carboxylate carbon of DHA and the terminal carbon of the side chains of residues near inner leaflet include R41, R42, R465, and K467. Different colored traces correspond to distinct DHA molecules. Only DHA molecules that approached within 5 Å of a given residue in at least one frame were included in the analysis.

**Supplementary Figure 3:**
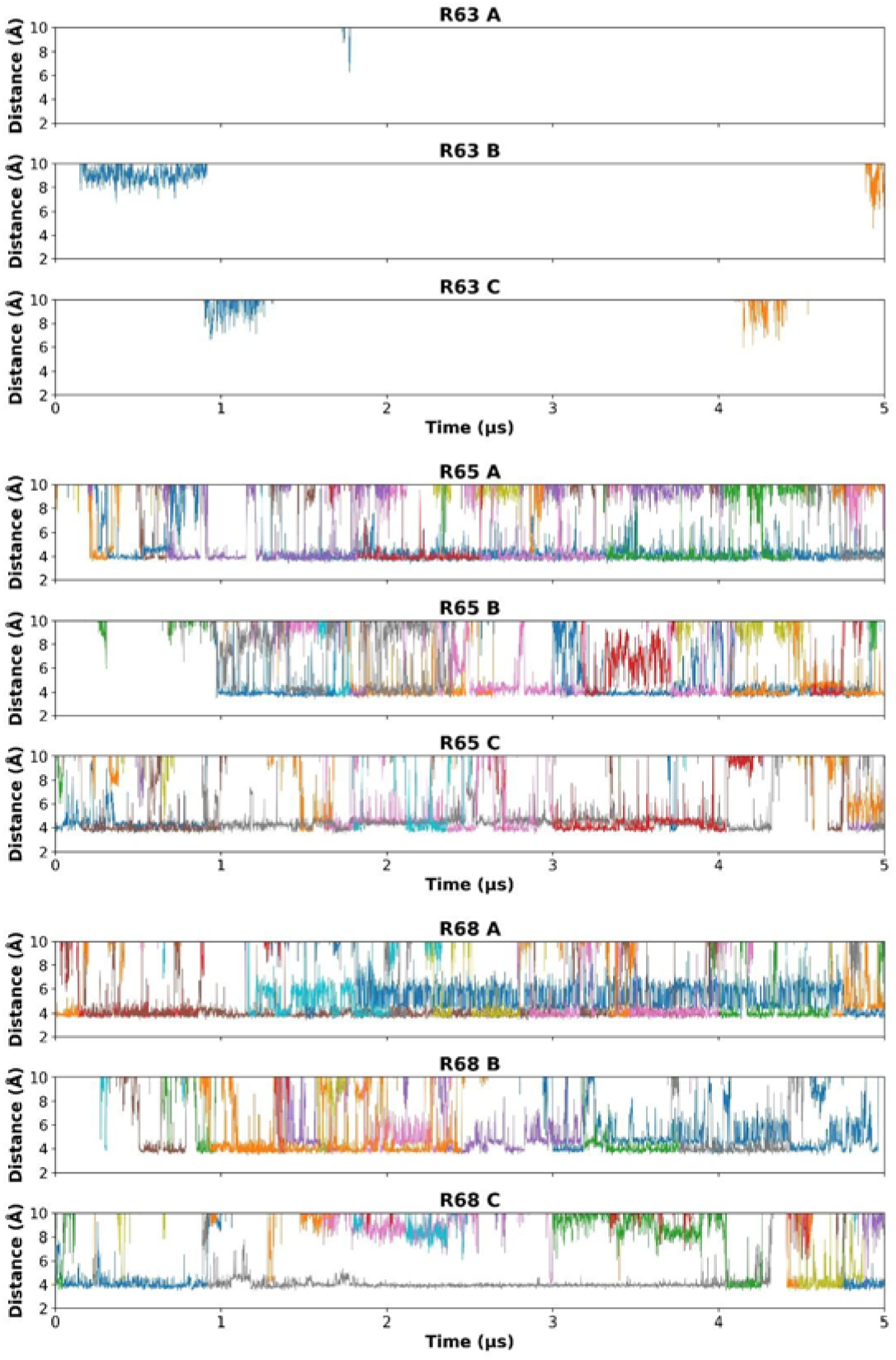
The spontaneous DHA binding to residues near outer leaflet in resting-state (Trajectory 1, the 5 µs simulation of the resting-state hASIC3 model with DHA randomly partitioned into the surrounding POPC bilayer; see Table 1): Distance plots showing the proximity between the carboxylate carbon of DHA and the terminal carbon of the side chains of arginine residues near outer leaflet (R63, R65 and R68) Different colored traces correspond to distinct DHA molecules. Only DHA molecules that approached within 5 Å of a given residue in at least one frame were included in the analysis.

**Supplementary Figure 4:**
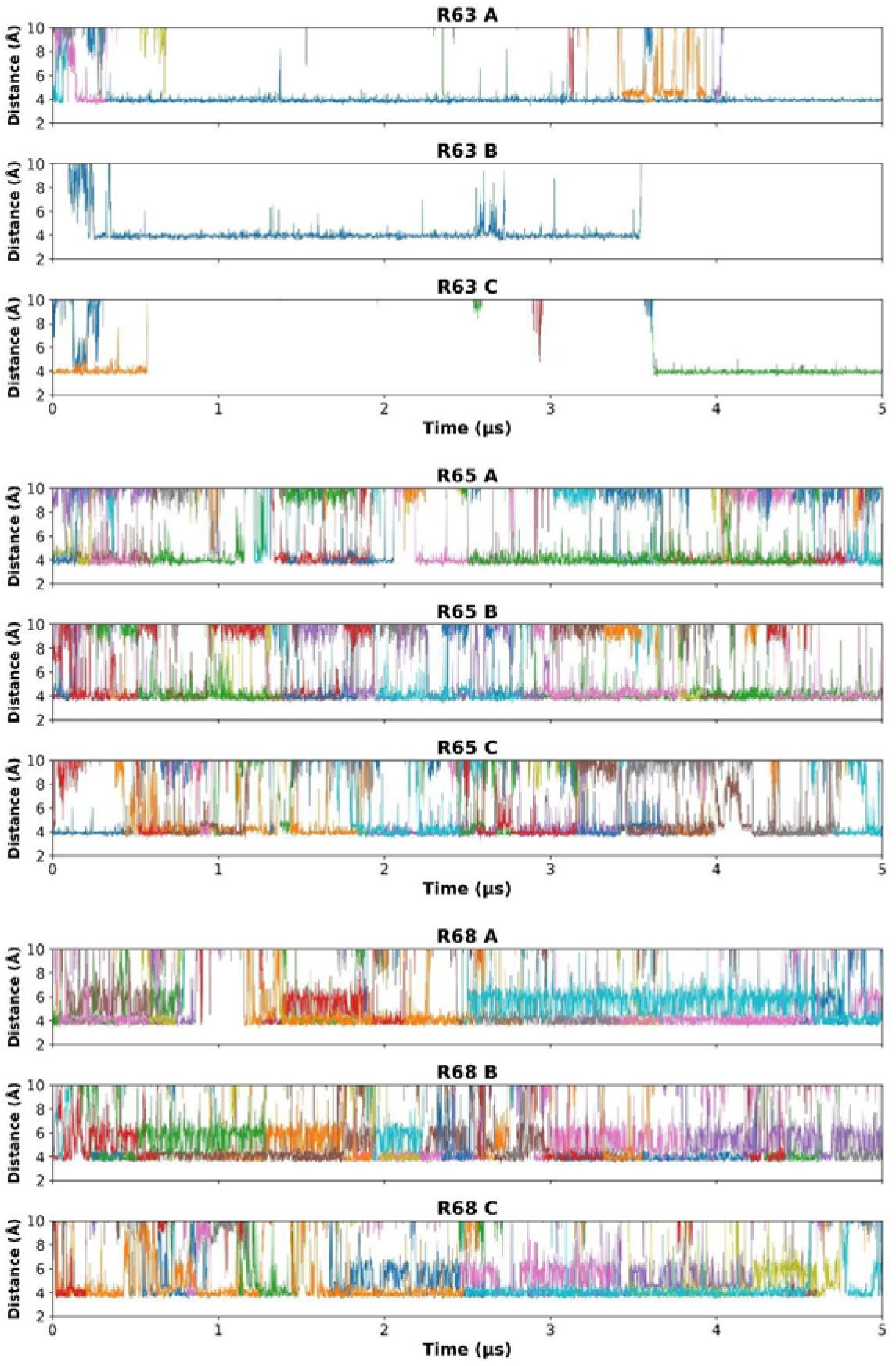
The spontaneous DHA binding in open state (Trajectory 2, the 5 µs simulation of the open-state hASIC3 model with DHA randomly partitioned into the surrounding POPC bilayer; see Table 1): Distance plots showing the proximity between the carboxylate carbon of DHA and the terminal carbon of the side chains of upper-leaflet residues R63, R65, and R68. Different colored traces correspond to distinct DHA molecules. Only DHA molecules that approached within 5 Å of a given residue in at least one frame were included in the analysis.

**Supplementary Figure 5:**
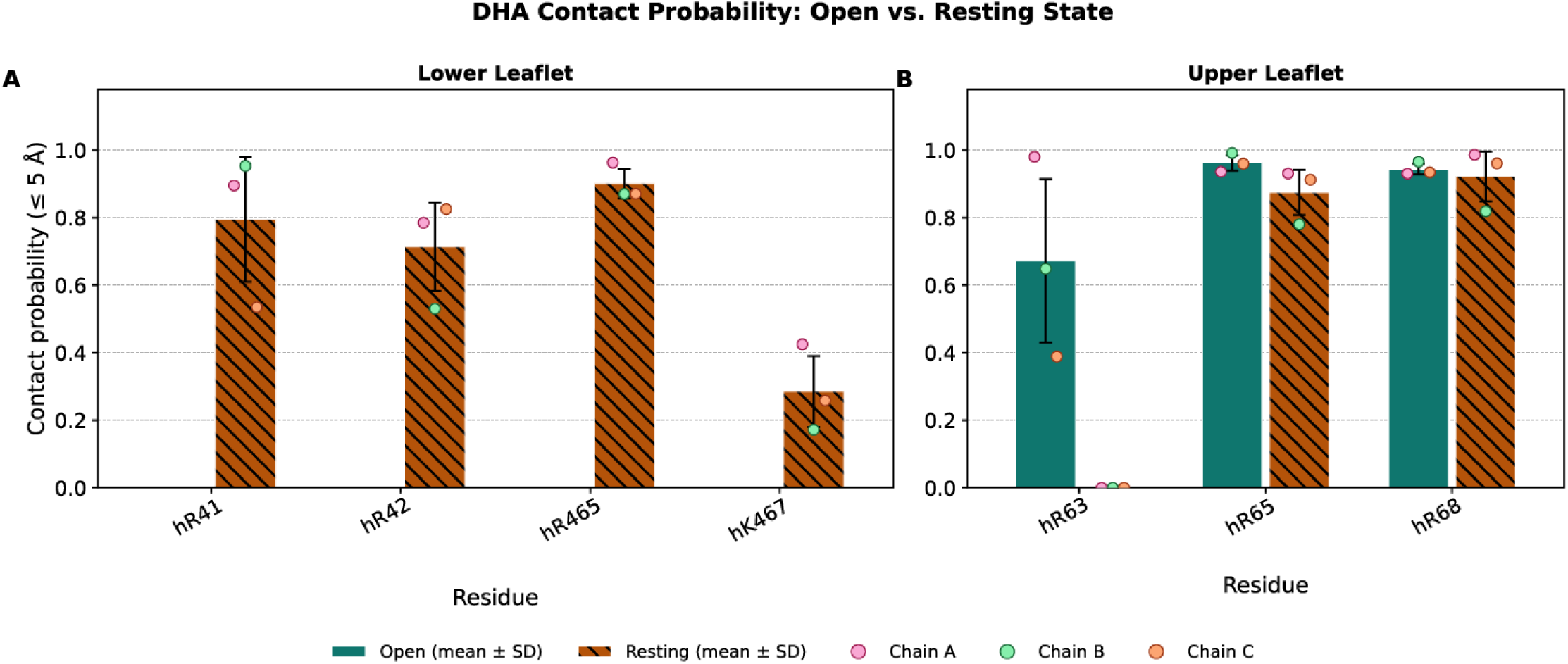
DHA contact probability with basic residues in open and resting states of ASIC3. Fraction of simulation frames in which at least one DHA molecule is within 5 Å of the indicated residue (minimum distance per frame; Trajectory 1, resting state, and Trajectory 2, open state; see Table 1). Bars represent mean ± SD across the three subunits, and points indicate individual chain values (Chains A–C). **(A)** Lower leaflet residues (hR41/rR42, hR42/rR43, hR465*, hK467/rR469). Data are shown for the resting state only, as these residues are not present in the open-state structural model. **(B)** Upper leaflet residues (hR63/rR64, hR65/rR66, hR68*) present in both open and resting state simulations. Asterisks denote residues without a conserved counterpart.

**Supplementary Figure 6.**
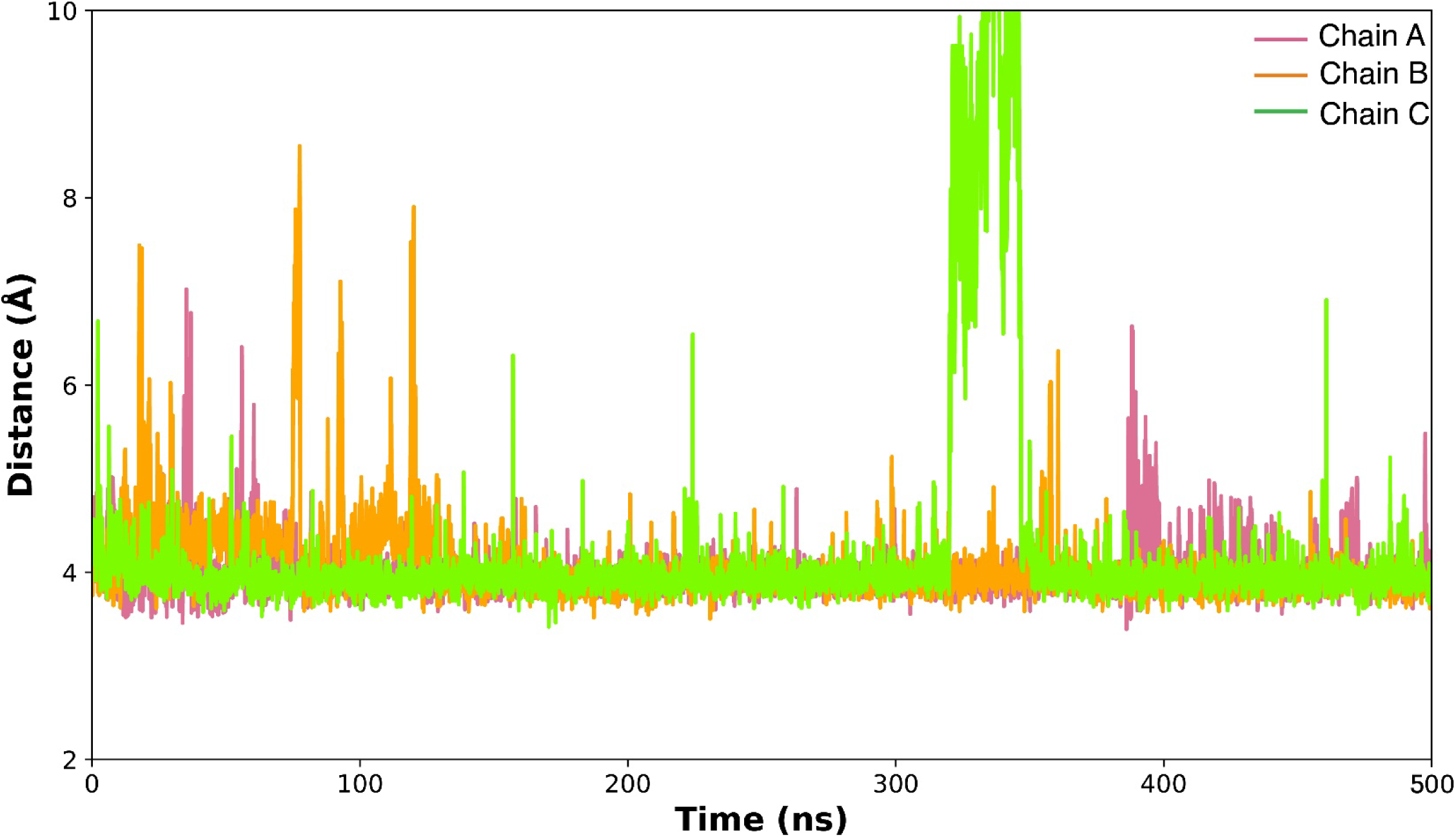
Stability of DHA binding to R63 across ASIC3 subunits. Time series of the distance between the DHA headgroup (carboxylate carbon) and the terminal carbon (Cζ) of the R63 side chain for each subunit (Chain A, pink; Chain B, orange; Chain C, green) over the course of the simulation (Trajectory 3, the 500 ns simulation of the open-state hASIC3 model with three DHA molecules pre-docked near the R63 binding site; see Table 1).

**Supplementary Figure 7:**
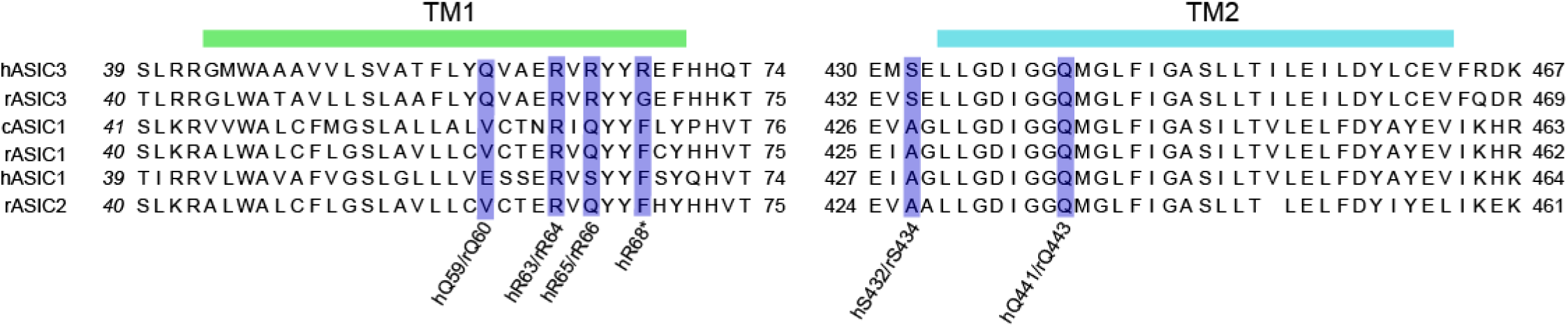
Sequence alignment for the segments TM1 and TM2 for rat ASIC1-3, human ASIC3, human ASIC1 and chicken ASIC1. Highlighted in purple are the residues that align with the critical DHA binding residues identified in rat and human ASIC3 across all ASIC genes.

**Supplementary Figure 8:**
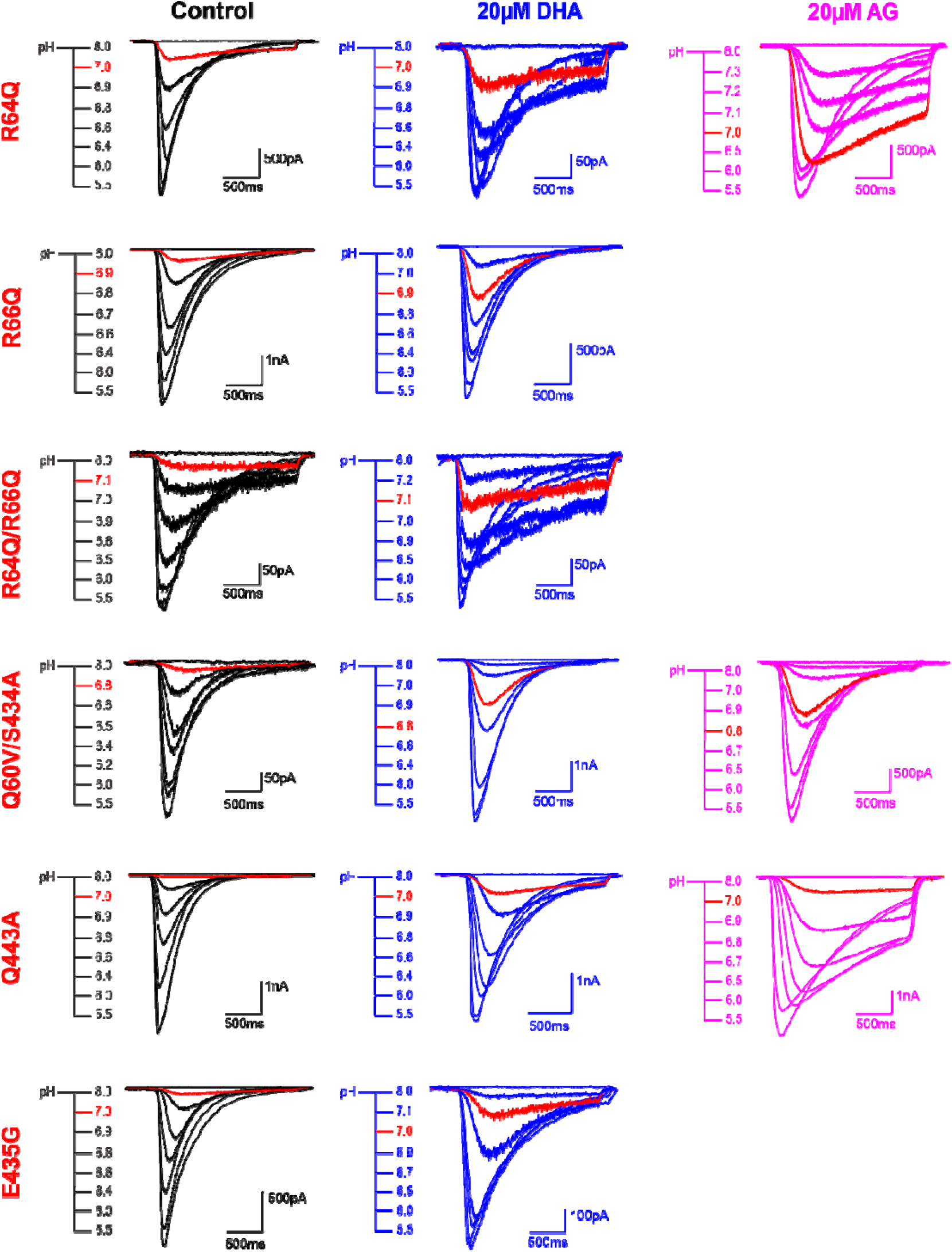
Representative traces of rat ASIC3 WT and mutants showing pH-dependent channel activation ± 20 µM DHA or AG application. Traces highlighted in red are to indicate the response to a specific pH across all conditions within each construct. For lipid conditions, cells were pre-incubated for at least 5 minutes prior to each experiment.

**Supplementary Table 1:** Decay rates (1/e) for rASIC3 WT and R64Q mutant channels ± 20 µM lipid applications. *n* denotes the number of individual cells. Statistical analysis was performed within each construct using a One-way ANOVA with Dunnett’s post-hoc multiple comparisons test. Mean differences and adjusted P values within each rASIC3 construct were obtained by comparing control vs. lipid conditions. WT and R64Q control and 20 µM DHA data from Figs. 1 and 6 respectively were used for comparison.

| rASIC3 Construct | Condition | 1/e (ms) | SEM | *n* | Mean diff. (Control vs. Lipid) | Adjusted P value |
| --- | --- | --- | --- | --- | --- | --- |
| WT | Control | 385.49 | 10.92 | 19 | -- | -- |
|  | 20μM ETA | 440.07 | 10.56 | 8 | -50.17 | 0.5774 |
|  | 20μM AA | 521.39 | 27.78 | 8 | -131.5 | 0.0038 |
|  | 20μM DHA | 545.83 | 30.76 | 17 | -160.3 | <0.0001 |
|  | 20μM AG | 636.87 | 31.90 | 9 | 247 | <0.0001 |
|  | 20μM AS | 799.96 | 30.016 | 11 | -410.1 | <0.0001 |
|  | 20μM LPC | 1107.44 | 37.63 | 8 | -717.5 | <0.0001 |
|  | 20μM AS | 799.96 | 30.016 | 11 | -410.1 | <0.0001 |
| R64Q | Control | 418.93 | 16.21 | 16 | -- | -- |
|  | 20μM AA | 410.67 | 16.33 | 8 | 8.253 | >0.9999 |
|  | 20μM AA | 410.67 | 16.33 | 8 | 8.253 | >0.9999 |
|  | 20μM DHA | 413.63 | 13.13 | 8 | 5.297 | >0.9999 |
|  | 20μM AG | 613.59 | 46.36 | 8 | -194.7 | 0.0018 |
|  | 20μM AS | 1078.05 | 58.06 | 11 | -659.1 | <0.0001 |
|  | 20μM LPC | 2084.75 | 143.55 | 7 | -1666 | <0.0001 |

**Supplementary Table 2:** Comparison of the magnitude of the effects of the lipids on each mutant. (top) Difference in the decay rates (1/e) for rASIC3 WT and mutant channels ± 20 µM lipid applications. *n* denotes the number of individual cells. A One-way ANOVA with Dunnett’s post-hoc multiple comparisons test was performed. (bottom) Difference in the pH_0.5_ for rASIC3 WT and mutant channels ± 20 µM lipid applications. A One-way ANOVA with Dunnett’s post-hoc multiple comparisons test was performed. Standard deviation (SD) for each is the result of propagating the error appropriately for the subtraction of two means. P values are the obtained by comparing the magnitude of the WT effect to each of the other mutants.

| rASIC3 Construct | Shift in pH<sub>0.5</sub> | SD | *n* | Adjusted P value |
| --- | --- | --- | --- | --- |
| WT | 0.24 | 0.06 | 9 | - |
| R64Q | 0.15 | 0.05 | 8 | 0.082 |
| R66Q | 0.14 | 0.07 | 8 | 0.037 |
| R64Q/R66Q | 0.18 | 0.05 | 11 | 0.312 |
| Q60V/S434A | 0.22 | 0.06 | 13 | 0.992 |
| Q443A | 0.18 | 0.09 | 10 | 0.395 |
| E435G | 0.14 | 0.06 | 11 | 0.025 |

**Supplementary Table 2:** Comparison of the magnitude of the effects of the lipids on each mutant.
| rASIC3 Construct | 1/e (ms) | SD | *n* | Adjusted P value |
| --- | --- | --- | --- | --- |
| WT | 160.35 | 131.46 | 17 | - |
| R64Q | 5.08 | 69.64 | 8 | 0.01 |
| R66Q | 83.71 | 88.49 | 10 | 0.481 |
| R64Q/R66Q | -8.77 | 127.26 | 10 | 0.001 |
| Q60V/S434A | 31.39 | 67.37 | 13 | 0.014 |
| Q443A | 99.32 | 92.03 | 11 | 0.704 |
| E435G | 162.53 | 83.00 | 10 | >0.9999 |

**Supplementary Table 3:** Decay rates (1/e) for ASIC3 WT and mutants ± 20 µM AG application. *n* denotes the number of individual cells. Statistical analysis was performed within each construct using unpaired t-tests with Welch’s correction. Mean differences and adjusted P values within each ASIC3 construct were obtained by comparing control vs. 20 µM AG. WT control data from Fig. 1 was used for comparison.

| rASIC3 Construct | Condition | 1/e (ms) | SEM | *n* | Mean diff. (Control vs. Lipid) | Adjusted P value |
| --- | --- | --- | --- | --- | --- | --- |
| WT | Control | 385.5 | 10.92 | 19 | -251.4 | <0.0001 |
| | 20μM AG | 636.9 | 31.90 | 9 | | |
| Q60V/S434A | Control | 318.0 | 16.10 | 13 | -151.0 | <0.0001 |
| | 20μM AG | 469.0 | 12.94 | 14 | | |
| Q443A | Control | 377.7 | 15.07 | 10 | -390.4 | <0.0001 |
| | 20μM AG | 768.1 | 36.92 | 7 | | |

**Supplementary Table 4:** Activation pH_0.5_ values for ASIC3 WT and mutants ± 20 µM AG application. *n* denotes the number of individual cells. Statistical analysis was performed within each construct using unpaired t-tests with Welch’s correction. Mean differences and adjusted P values within each ASIC3 construct were obtained by comparing control vs. 20 µM AG. WT control data from Fig. 1 was used for comparison.

| rASIC3 Construct | Condition | pH<sub>0.5</sub> | SEM | *n* | Mean diff. (Control vs. Lipid) | Adjusted P value |
| --- | --- | --- | --- | --- | --- | --- |
| **WT** | Control | 6.606 | 0.014 | 13 | -0.2562 | <0.0001 |
|  | 20μM AG | 6.862 | 0.027 | 5 |  |  |
| **R64Q** | Control | 6.866 | 0.012 | 13 | -0.2625 | <0.0001 |
|  | 20μM AG | 7.129 | 0.028 | 8 |  |  |
| **Q60V/S434A** | Control | 6.415 | 0.012 | 13 | -0.2113 | <0.0001 |
|  | 20μM AG | 6.627 | 0.010 | 13 |  |  |
| **Q443A** | Control | 6.617 | 0.025 | 9 | -0.1507 | 0.0049 |
|  | 20μM AG | 6.768 | 0.026 | 6 |  |  |

## References

1. Zaydman, M. A. & Cui, J. PIP2 regulation of KCNQ channels: biophysical and molecular mechanisms for lipid modulation of voltage-dependent gating. Front. Physiol. 5, (2014).

2. Sun, J. & MacKinnon, R. Structural Basis of Human KCNQ1 Modulation and Gating. Cell 180, 340–347.e9 (2020).

3. Klipp, R. C. & Bankston, J. R. Structural determinants of acid-sensing ion channel potentiation by single chain lipids. Journal of General Physiology 154, e202213156 (2022).

4. Mathiharan, Y. K. et al. Structural insights into GIRK2 channel modulation by cholesterol and PIP2. Cell Reports 36, 109619 (2021).

5. Handlin, L. J. & Dai, G. Direct regulation of the voltage sensor of HCN channels by membrane lipid compartmentalization. Nat Commun 14, 6595 (2023).

6. Tian, Y. et al. Atomic determinants of BK channel activation by polyunsaturated fatty acids. PNAS 113, 13905–13910 (2016).

7. Levental, I. & Lyman, E. Regulation of membrane protein structure and function by their lipid nano-environment. Nat Rev Mol Cell Biol 24, 107–122 (2023).

8. Lemmon, M. A. Membrane recognition by phospholipid-binding domains. Nat Rev Mol Cell Biol 9, 99– 111 (2008).

9. Phillips, R., Ursell, T., Wiggins, P. & Sens, P. Emerging roles for lipids in shaping membrane-protein function. Nature 459, 379–385 (2009).

10. Tong, A. et al. Direct binding of phosphatidylglycerol at specific sites modulates desensitization of a ligand-gated ion channel. eLife 8, e50766 (2019).

11. Bergh, C., Rovšnik, U., Howard, R. & Lindahl, E. Discovery of lipid binding sites in a ligand-gated ion channel by integrating simulations and cryo-EM. eLife 12, RP86016 (2024).

12. Zheng, H., Liu, W., Anderson, L. Y. & Jiang, Q.-X. Lipid-dependent gating of a voltage-gated potassium channel. Nat Commun 2, 250 (2011).

13. Gonen, T., Sliz, P., Kistler, J., Cheng, Y. & Walz, T. Aquaporin-0 membrane junctions reveal the structure of a closed water pore. Nature 429, 193–197 (2004).

14. Marra, S. et al. Non-acidic activation of pain-related Acid-Sensing Ion Channel 3 by lipids. EMBO J 35, 414–428 (2016).

15. Smith, E. S., Cadiou, H. & McNaughton, P. A. Arachidonic acid potentiates acid-sensing ion channels in rat sensory neurons by a direct action. Neuroscience 145, 686–698 (2007).

16. Allen, N. J. & Attwell, D. Modulation of ASIC channels in rat cerebellar purkinje neurons by ischaemia-related signals. J Physiol 543, 521–529 (2002).

17. Heusser, S. A. & Pless, S. A. Acid-sensing ion channels as potential therapeutic targets. Trends in Pharmacological Sciences 42, 1035–1050 (2021).

18. Boscardin, E., Alijevic, O., Hummler, E., Frateschi, S. & Kellenberger, S. The function and regulation of acid-sensing ion channels (ASICs) and the epithelial Na(+) channel (ENaC): IUPHAR Review 19. Br J Pharmacol 173, 2671–2701 (2016).

19. Wood, J. N. II. Genetic approaches to pain therapy. American Journal of Physiology-Gastrointestinal and Liver Physiology 278, G507–G512 (2000).

20. Poirot, O., Berta, T., Decosterd, I. & Kellenberger, S. Distinct ASIC currents are expressed in rat putative nociceptors and are modulated by nerve injury. The Journal of Physiology 576, 215–234 (2006).

21. White, J. P. M. et al. Role of transient receptor potential and acid-sensing ion channels in peripheral inflammatory pain. Anesthesiology 112, 729–741 (2010).

22. Deval, E. et al. ASIC3, a sensor of acidic and primary inflammatory pain. The EMBO Journal 27, 3047– 3055 (2008).

23. Ikeuchi, M., Kolker, S. J., Burnes, L. A., Walder, R. Y. & Sluka, K. A. Role of ASIC3 in the primary and secondary hyperalgesia produced by joint inflammation in mice. PAIN® 137, 662–669 (2008).

24. Mamet, J., Baron, A., Lazdunski, M. & Voilley, N. ProInflammatory Mediators, Stimulators of Sensory Neuron Excitability via the Expression of Acid-Sensing Ion Channels. J. Neurosci. 22, 10662–10670 (2002).

25. Fraser, D. D. et al. Elevated polyunsaturated fatty acids in blood serum obtained from children on the ketogenic diet. Neurology 60, 1026–1029 (2003).

26. Hammarström, S. et al. Increased concentrations of nonesterified arachidonic acid, 12L-hydroxy-5,8,10,14-eicosatetraenoic acid, prostaglandin E2, and prostaglandin F2alpha in epidermis of psoriasis. Proceedings of the National Academy of Sciences 72, 5130–5134 (1975).

27. Siddiqui, R. A., Harvey, K. A. & Zaloga, G. P. Modulation of enzymatic activities by n-3 polyunsaturated fatty acids to support cardiovascular health. The Journal of Nutritional Biochemistry 19, 417–437 (2008).

28. Wiktorowska-Owczarek, A., Berezińska, M. & Nowak, J. Z. PUFAs: Structures, Metabolism and Functions. Adv Clin Exp Med 24, 931–941 (2015).

29. Ananchenko, A. & Musgaard, M. Multiscale molecular dynamics simulations predict arachidonic acid binding sites in human ASIC1a and ASIC3 transmembrane domains. Journal of General Physiology 155, e202213259 (2023).

30. Jowais, J. J. et al. Mechanistic understanding of KCNQ1 activating polyunsaturated fatty acid analogs. J Gen Physiol 155, e202313339 (2023).

31. Yazdi, S. et al. Identification of PUFA interaction sites on the cardiac potassium channel KCNQ1. J Gen Physiol 153, e202012850 (2021).

32. Sun, D. et al. Structural insights into human acid-sensing ion channel 1a inhibition by snake toxin mambalgin1. eLife 9, e57096 (2020).

33. Baconguis, I., Bohlen, C. J., Goehring, A., Julius, D. & Gouaux, E. X-Ray Structure of Acid-Sensing Ion Channel 1–Snake Toxin Complex Reveals Open State of a Na+-Selective Channel. Cell 156, 717–729 (2014).

34. Jacquot, F. et al. Lysophosphatidylcholine 16:0 mediates chronic joint pain associated to rheumatic diseases through acid-sensing ion channel 3. Pain 163, 1999–2013 (2022).

35. Kundu, A., Yamaguchi, S. & Tahara, T. Evaluation of pH at Charged Lipid/Water Interfaces by Heterodyne-Detected Electronic Sum Frequency Generation. J. Phys. Chem. Lett. 5, 762–766 (2014).

36. Kundu, A., Yamaguchi, S. & Tahara, T. Local pH at Nonionic and Zwitterionic Lipid/Water Interfaces Revealed by Heterodyne-Detected Electronic Sum-Frequency Generation: A Unified View to Predict Interfacial pH of Biomembranes. J. Phys. Chem. B 127, 5445–5452 (2023).

37. Romero, L. O. et al. Dietary fatty acids fine-tune Piezo1 mechanical response. Nat Commun 10, 1200 (2019).

38. Martinière, A. et al. Uncovering pH at both sides of the root plasma membrane interface using noninvasive imaging. Proceedings of the National Academy of Sciences 115, 6488–6493 (2018).

39. Klipp, R. C., Cullinan, M. M. & Bankston, J. R. Insights into the molecular mechanisms underlying the inhibition of acid-sensing ion channel 3 gating by stomatin. Journal of General Physiology 152, e201912471 (2020).

40. Bandarupalli, R., Roth, R., Klipp, R. C., Bankston, J. R. & Li, J. Molecular Insights into Single-Chain Lipid Modulation of Acid-Sensing Ion Channel 3. J. Phys. Chem. B 128, 12685–12697 (2024).

41. Waterhouse, A. et al. SWISS-MODEL: homology modelling of protein structures and complexes. Nucleic Acids Res 46, W296–W303 (2018).

42. Schrödinger Release 2025-4. Schrödinger, LLC (2025).

43. Lee, J. et al. CHARMM-GUI Input Generator for NAMD, GROMACS, AMBER, OpenMM, and CHARMM/OpenMM Simulations Using the CHARMM36 Additive Force Field. J. Chem. Theory Comput. 12, 405–413 (2016).

44. Jo, S., Kim, T., Iyer, V. G. & Im, W. CHARMM-GUI: a web-based graphical user interface for CHARMM. J Comput Chem 29, 1859–1865 (2008).

45. Olsson, M. H. M., Søndergaard, C. R., Rostkowski, M. & Jensen, J. H. PROPKA3: Consistent Treatment of Internal and Surface Residues in Empirical pKa Predictions. J Chem Theory Comput 7, 525–537 (2011).

46. Lomize, M. A., Pogozheva, I. D., Joo, H., Mosberg, H. I. & Lomize, A. L. OPM database and PPM web server: resources for positioning of proteins in membranes. Nucleic Acids Res 40, D370–D376 (2012).

47. Phillips, J. C. et al. Scalable molecular dynamics with NAMD. J Comput Chem 26, 1781–1802 (2005).

48. Shaw, D. E. et al. Anton 2: Raising the Bar for Performance and Programmability in a Special-Purpose Molecular Dynamics Supercomputer. in SC ’14: Proceedings of the International Conference for High Performance Computing, Networking, Storage and Analysis 41–53 (2014). doi:10.1109/SC.2014.9.

49. CHARMM36 all-atom additive protein force field: Validation based on comparison to NMR data – Huang - 2013 - Journal of Computational Chemistry - Wiley Online Library. https://onlinelibrary.wiley.com/doi/10.1002/jcc.23354.

50. Huang, J. et al. CHARMM36m: an improved force field for folded and intrinsically disordered proteins. Nat Methods 14, 71–73 (2017).

51. MacKerell, A. D., Banavali, N. & Foloppe, N. Development and current status of the CHARMM force field for nucleic acids. Biopolymers 56, 257–265 (2000).

52. Best, R. B. et al. Optimization of the Additive CHARMM All-Atom Protein Force Field Targeting Improved Sampling of the Backbone P, ψ and Side-Chain χ1 and χ2 Dihedral Angles. J. Chem. Theory Comput. 8, 3257–3273 (2012).

53. Klauda, J. B. et al. Update of the CHARMM All-Atom Additive Force Field for Lipids: Validation on Six Lipid Types. J. Phys. Chem. B 114, 7830–7843 (2010).

54. Jorgensen, W. L., Chandrasekhar, J., Madura, J. D., Impey, R. W. & Klein, M. L. Comparison of simple potential functions for simulating liquid water. J. Chem. Phys. 79, 926–935 (1983).

55. Darden, T., York, D. & Pedersen, L. Particle mesh Ewald: An N⋅log(N) method for Ewald sums in large systems. J. Chem. Phys. 98, 10089–10092 (1993).

56. Ryckaert, J.-P., Ciccotti, G. & Berendsen, H. J. C. Numerical integration of the cartesian equations of motion of a system with constraints: molecular dynamics of n-alkanes. Journal of Computational Physics 23, 327–341 (1977).

57. Hopkins, C. W., Le Grand, S., Walker, R. C. & Roitberg, A. E. Long-Time-Step Molecular Dynamics through Hydrogen Mass Repartitioning. J. Chem. Theory Comput. 11, 1864–1874 (2015).

58. Shan, Y., Klepeis, J. L., Eastwood, M. P., Dror, R. O. & Shaw, D. E. Gaussian split Ewald: A fast Ewald mesh method for molecular simulation. J Chem Phys 122, 54101 (2005).

59. Humphrey, W., Dalke, A. & Schulten, K. VMD: Visual molecular dynamics. Journal of Molecular Graphics 14, 33–38 (1996).

